# Timing-dependent modulation of working memory by VTA dopamine release in medial prefrontal cortex

**DOI:** 10.1101/2023.09.01.555676

**Authors:** Chaofan Ge, Zhaoqin Chen, Fangmiao Sun, Ruiqing Hou, Hongmei Fan, Yulong Li, Chengyu T. Li

**Affiliations:** Institute of Neuroscience, Center for Excellence in Brain Science and Intelligence Technology, Chinese Academy of Sciences, State Key Laboratory of Neuroscience, Shanghai, China; Shanghai Center for Brain Science and Brain-Inspired Technology, Shanghai, China; Peking University, Beijing, China; Lingang Laboratory, Shanghai, China

**Author notes:** One sentence summary: Dopamine in the medial prefrontal cortex (mPFC) modulates working memory in a timing-dependent and bidirectional manner through modifying activity gain and memory-coding ability of the mPFC neurons, as revealed by sensor imaging, pathway-specific optogenetics and electrophysiological recordings.

**Keywords:** Keyword Working memory, dopamine, medial prefrontal cortex, bidirectional, timing-dependent, activity gain, memory encoding, ventral tegmental area, optogenetics

## Abstract

Dopamine significantly modulates working memory (WM)^1–8^, a fundamental cognitive function that maintains information during a brief delay period^9–11^. The temporal precision of dopaminergic modulation in WM and related neural mechanisms remain elusive. Here we unveiled the pivotal role of dopamine timing in WM performance with head-fixed mice engaged in learning an olfactory WM task. Through electrophysiology and optogenetics, we found that dopaminergic neurons in the ventral tegmental area (VTA) could encode WM information during the delay period, and manipulating dopaminergic neuronal activity bidirectionally modulated WM performance in an inverted U-shaped manner. Optogenetic manipulation of VTA activity also induced bidirectional changes in WM-related neural activity in the medial prefrontal cortex (mPFC). Imaging with a dopamine-sensitive GPCR-activation-based sensor (GRAB_DA_)^12,13^ revealed transient dopamine peaks in the mPFC specifically during the early-delay period. Optogenetic stimulation of VTA-to-mPFC dopaminergic projections during the early- and late-delay period enhanced and impaired WM performance, respectively. Manipulations outside these periods or involving the nucleus accumbens had no effect. Single-unit recordings demonstrated that optogenetic excitation of VTA-to-mPFC projections could modulate mPFC neuronal activity in a manner consistent with behavioral modulation. Thus, the timing of dopamine release is critical in modulation of neuronal activity and behavior.

## INTRODUCTION

Dopamine in ventral tegmental area (VTA) plays crucial roles in modulating cognition and behavior^14–16^, including reward^16,17^, sensory perception^18–21^, motor action^22^, timing estimation^23,24^, etc.. VTA dopamine neurons could encode differential external and internal information at various behavioral phases^25^. In particular, dopamine plays crucial roles in working memory (WM)^1–8^, the process of actively maintaining and manipulating information for a brief delay period of several seconds^9–11^. Midbrain dopaminergic neurons in monkeys are typically activated during the sensory-delivery period, with small but significant elevation during the delay period^26^. Optogenetic tagging experiment in mice revealed that VTA dopaminergic neurons could encoding spatial WM information^27^.

The prefrontal cortex (PFC) is critical for WM^11,28^ and is innerved by dopaminergic terminals^29^. Pharmacological manipulation of DA receptors in PFC could modulate WM related neuronal activity and performance in monkeys^5,6,30–33,31^ and rodents^34^ performing WM tasks. Temporal dynamics of dopaminergic activity in PFC play important roles in various behavioral tasks, as shown by the results that phasic and tonic activation of VTA projections to PFC induce differential behavioral modulations^35,36^.

However, the causal importance of temporal dynamics in dopamine release for WM-task performance and its neural mechanism remain unclear. In this study, we combined sensor imaging, optogenetic manipulation, and electrophysiological recording to reveal the importance of dopamine timing in modulating WM behavior and its neural mechanisms.

## Results

### Importance of VTA dopaminergic neurons in WM performance and information encoding

We trained head-fixed mice to perform an olfactory delayed paired-association (ODPA) task (Fig.1a; as in Ref.37-39). In each trial, we delivered a sample odor (S1 or S2), then following a 5- or 6-sec delay period, a test odor (T1 or T2). Mice were rewarded with water if they licked within a response window in the defined “paired” trials (S1-T1 or S2-T2), but not in the “unpaired” trials (S1-T2 or S2-T1). Sample-odor information needs to be maintained in WM for efficient task performance. In several days, mice learned to withhold from licking during the response window in the unpaired trials (Extended Data Fig.1a). They showed little licking during the delay period (Extended Data Fig.1b).

**Fig.1.**
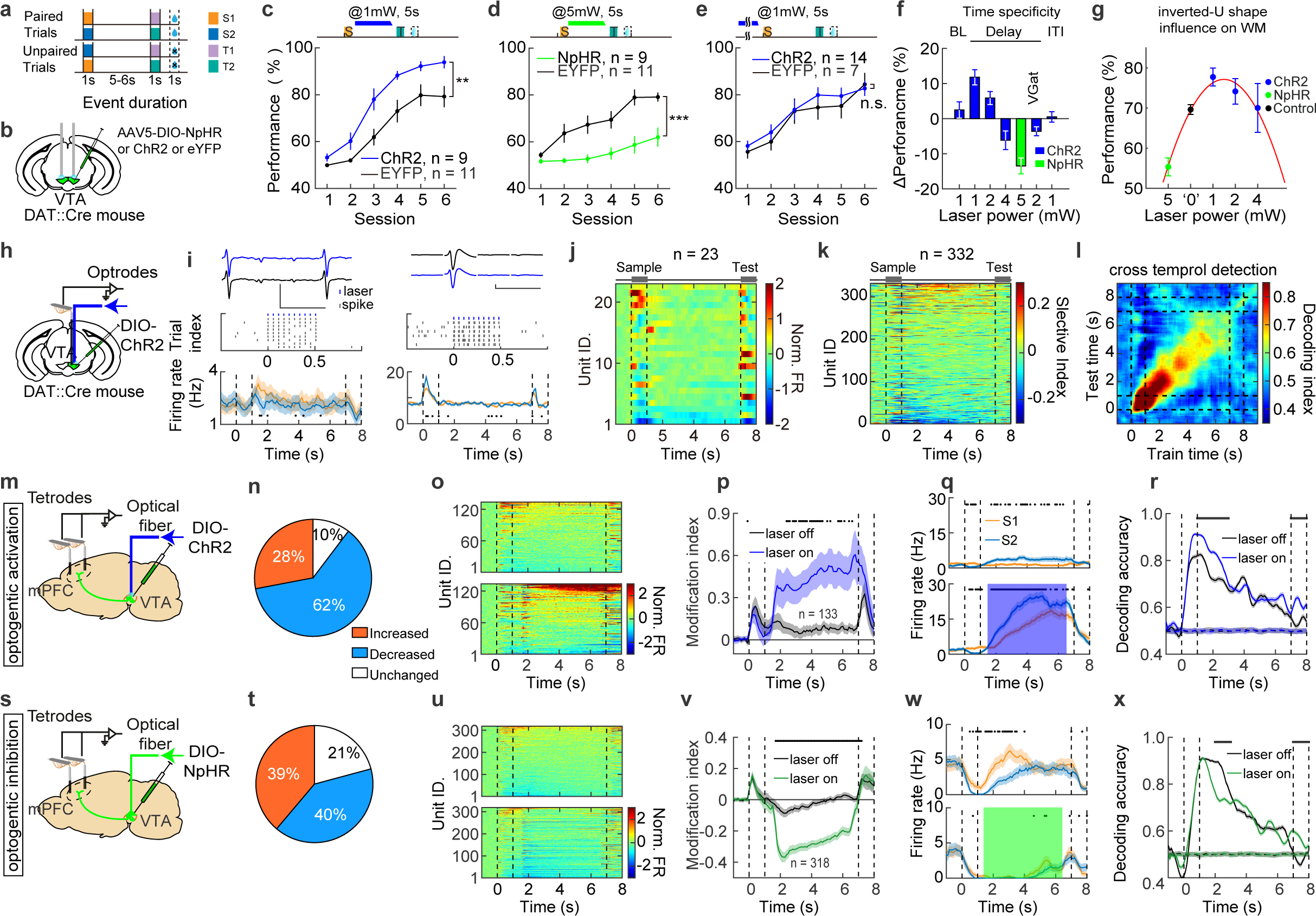
Dopamine neurons in VTA bilaterally modulated WM and memory-coding ability of mPFC neurons. **a**, Schematic diagram of the olfactory delayed paired association (ODPA) task. **b**, Strategy for optogenetic activation or inhibition of VTA^DA^ neurons. **c**, Behavioral performance with optogenetic activation of VTA^DA^ neurons during the delay period. Error bars: standard error of the mean (SEM) unless stated otherwise. **p = 0.0026, Two-way ANOVA with repeated measure. **d**, Performance with optogenetic inhibition of VTA^DA^ neurons during the delay period. *** p < 0.001, Two-way ANOVA with repeated measure. **e**, Performance with optogenetic activation of VTA^DA^ neurons during the baseline period. (N.S., p > 0.05, Two-way ANOVA with repeated measure). **f**, Temporal specificity in optogenetic activation and inhibition of dopamine neurons, showing averaged changes in performance of the 6 training days. Data from **c-e** and Extended data Fig.2 averaged cross 6 sessions. **g**, Inverted-U shape in performance following optogenetic manipulation of VTA^DA^ neurons during the delay period. Data from **c-d** and Extended data Fig.2d,g (n = 9 for NpHR, n = 44 for control, n = 9 for ChR2 mice with 1mW laser, n = 9 for ChR2 mice with 2 mW laser, and n = 7 for ChR2 mice with 4 mW laser, data were averaged cross 6 sessions for each mouse). **h**, Strategy for electrophysiological recoding of VTA^DA^ neurons. **i**, an example opto-tagged VTA^DA^ neuron. Above: waveforms recorded from the tetrode (blue for laser triggered waveforms and black for spontaneous ones). Scale bars for waveforms plots throughout all figures: x axis, 2 ms; y axis, 100 μV. Middle: raster showing action potentials following laser stimuli. The blue ticks indicate laser pulses and the grey ticks indicate the spikes. Below: Peri-stimulus time histogram (PSTH) showing activity of the neuron given S1 (orange) and S2 (blue) as sample odors. Black dots below indicate bins with statistically significant difference (p < 0.05, Wilcoxon rank sum test, 100-ms bin) for all figures, unless stated otherwise. Surrounding shades represent SEM for all figures, unless stated otherwise. **j**, Another example opto-tagged VTA^DA^ neurons. **k**, Heatmap showing neural modulation during ODPA task for 23 tagged VTA^DA^ neurons. **l**, Heatmap showing odor-selectivity index for the selective units in VTA (n = 332 from 1251 units). **m**, Cross-temporal decoding result for the units in **l**. **n**, Strategy for optogenetic excitation of VTA^DA^ neurons during the delay period while simultaneously recording mPFC neuronal activity. **o**, Proportion of activity modulation in mPFC neurons following optogenetic excitation of VTA^DA^ neurons (Increased: n = 133; Decreased: n = 293; Unchanged: n = 49). **p**, Activity heatmaps showing the mPFC units with enhanced activity in the laser-on (below) trials comparing to the laser-off (above) trials (n = 133). **q**, PSTH of normalized firing for the neurons in **p**. **r**, An example mPFC neuron showing increased sample selectivity during the delay period with activation of VTA^DA^ neurons. Above: laser off. Below: laser on. **s**, SVM decoding accuracy for sample odors, calculated by the mPFC neurons with enhanced firing following activation of VTA^DA^ neurons. **t**, Strategy for optogenetic suppression of VTA^DA^ neurons during the delay period while simultaneously recording mPFC neuronal activity. **u**, Proportion of activity modulation in mPFC neurons following suppression of VTA^DA^ neurons (Increased: n = 306; Decreased: n = 318; Unchanged: n = 169). **v**, Activity heatmaps showing the mPFC units with decreased activity in the laser-on (below) trials comparing to the laser-off (above) trials (n = 318). **q**, PSTH of normalized firing for the neurons in **v**. **x**, an example mPFC neuron showing reduced sample selectivity during the delay period with optogenetic suppression of VTA^DA^ neurons. Above: laser off. Below: laser on **y**, SVM decoding accuracy of mPFC neurons with decreased activity following optogenetic suppression of VTA^DA^ neurons.

We utilized cell type-specific optogenetic manipulation to either excite or suppress activity of VTA dopaminergic (VTA^DA^) neurons. AAV-DIO-ChR2 was injected in VAT of DAT-Cre mice to expression ChR2 in dopaminergic neurons (Fig.1b, Extended Data Fig.1c). In the learning phase, weak excitation (1 mw, blue laser of 473nm) during the delay period significantly improved WM performance (Fig.1c, Extended Data Fig.1e,h), mild excitation (2 mW) did not significantly modulate performance (Extended Data Fig.2d-f), whereas strong excitation (4 mW) impaired performance (Extended Data Fig.2g-i). Optogenetic excitation during either baseline (Fig.1e, Extended Data Fig.1f,i) or inter-trial interval (Extended Data Fig.2a-c) did not modulate performance.

To suppress delay-period activity of dopaminergic neurons, we injected AAV-DIO-NpHR into VTA of DAT-Cre mice (Fig.1b, Extended Data Fig.1c). Optogenetic suppression of VTA^DA^ neurons during the delay period impaired WM performance in the learning phase (green laser of 532nm; Fig.1d, Extended Data Fig.1d,g, Extended Data Fig.2j-k). Thus, delay-period activity of VTA^DA^ neurons bidirectionally modulated WM performance in an inverted U-shape manner (Fig.1f,g), consistent with previous reports^40–43^.

We then examined the population activation of VTA^DA^ neurons by fiber photometry, with AAV-DIO-GCaMP6 injected into VTA of DAT-Cre mice (Extended Data Fig.3a). Calcium imaging in VTA^DA^ neurons revealed a stronger activation during the test odor-delivery and reward periods, and a smaller yet significant peak during the sample odor-delivery period (Extended Data Fig.3).

Fiber-photometry imaging does not have single-cell resolution. We therefore examined the coding ability of VTA^DA^ neurons by tetrode recordings (Fig.1h, Extended Data Fig.4a), with optogenetic tagging (injecting AAV-DIO-ChR2 in VTA of DAT-Cre mice, two example opto-tagged neuron in Fig.1i-j; all tagged neurons in Fig.1k). Both tagged neurons (Fig.1i-j) and some non-tagged neurons (Extended data Fig.4b) showed characteristics of dopaminergic neurons, including strong positive and phasic responses to reward^16^ (Extended data Fig.4c). Overall, we observed differential activity following sample odors in many of the recorded VTA neurons (Fig.1l, Extended data Fig.4d for tagged VTA^DA^ neurons). Cross-temporal decoding (CTD) analysis revealed transient patterns of decoding for sample-odor information in VTA neurons, with significant coding ability along diagonal CTD and higher decoding ability during the early-delay period (Fig.1m, CTD was based on the template matching method). Thus, VTA^DA^ neurons showed coding ability of WM information, especially during the early-delay period (Extended Data Fig.4f).

To explore the cortical mechanisms underlying dopamine modulation in WM performance, we recorded the neuronal activity of mPFC while manipulating the delay-period activity of VTA^DA^ neurons. Tetrodes were inserted in mPFC and AAV-DIO-ChR2 was injected in VTA of DAT-Cre mice (Fig.1n, Extended Data Fig.5a). Optogenetic excitation by blue laser (473nm) during the delay period significantly modified neuronal activity, with 28% and 62% mPFC neurons increased and decreased their activity, respectively (Fig.1o, example neurons in Extended Data Fig.5b-g). Heatmap and averaged firing rate of mPFC neurons with elevated activity are shown in Fig.1p and 1q. We observed overall enhanced coding ability of WM information in the neurons with elevated activity upon dopaminergic excitation (example neuron in Fig.1r, overall decoding in Fig.1s). The activity modulation and coding ability of the mPFC neurons with reduced activity upon dopaminergic excitation are shown in Extended Data Fig.5.

Optogenetic suppression and recording mPFC activity were achieved by inserting tetrodes in mPFC and expressing AAV-DIO-NpHR in VTA of DAT-Cre mice (Fig.1t and Extended Data Fig.6a). Optogenetic suppression by green laser during the delay period strongly modified neuronal activity, with 39% and 40% mPFC neurons increased and decreased their activity, respectively (Fig.1u, example neurons in Extended Data Fig.6b-g). Heat map and averaged firing rate of mPFC neurons with reduced activity are shown in Fig.1v and 1w. We observed overall impaired coding ability of WM information in the neurons with reduced activity upon dopaminergic suppression (example neuron in Fig.1x, overall decoding in Fig.1y). The activity modulation and coding ability of the mPFC neurons with elevated activity upon dopaminergic suppression are shown in Extended Data Fig.6. Therefore, mPFC neuronal activity was significantly modified by optogenetic manipulation of VTA^DA^ neurons.

### Transient dopamine dynamics in mPFC associated with WM

Dopamine release in the mPFC has been observed with indirect and slow methodologies in performing WM tasks^43–45^. Here we monitored endogenous dopamine dynamics at a higher temporal resolution using a genetically-encoded GPCR-Activation-Based-DA (GRAB_DA_) sensor^13^ (Fig.2a-b). We expressed the virus of AAV-hSyn-GRABDA2m in the mPFC, and implanted optical fibers on top of the virus-injected area. After several weeks of viral expression, fiber-photometry imaging was performed while mice were learning the ODPA task.

**Fig.2.**
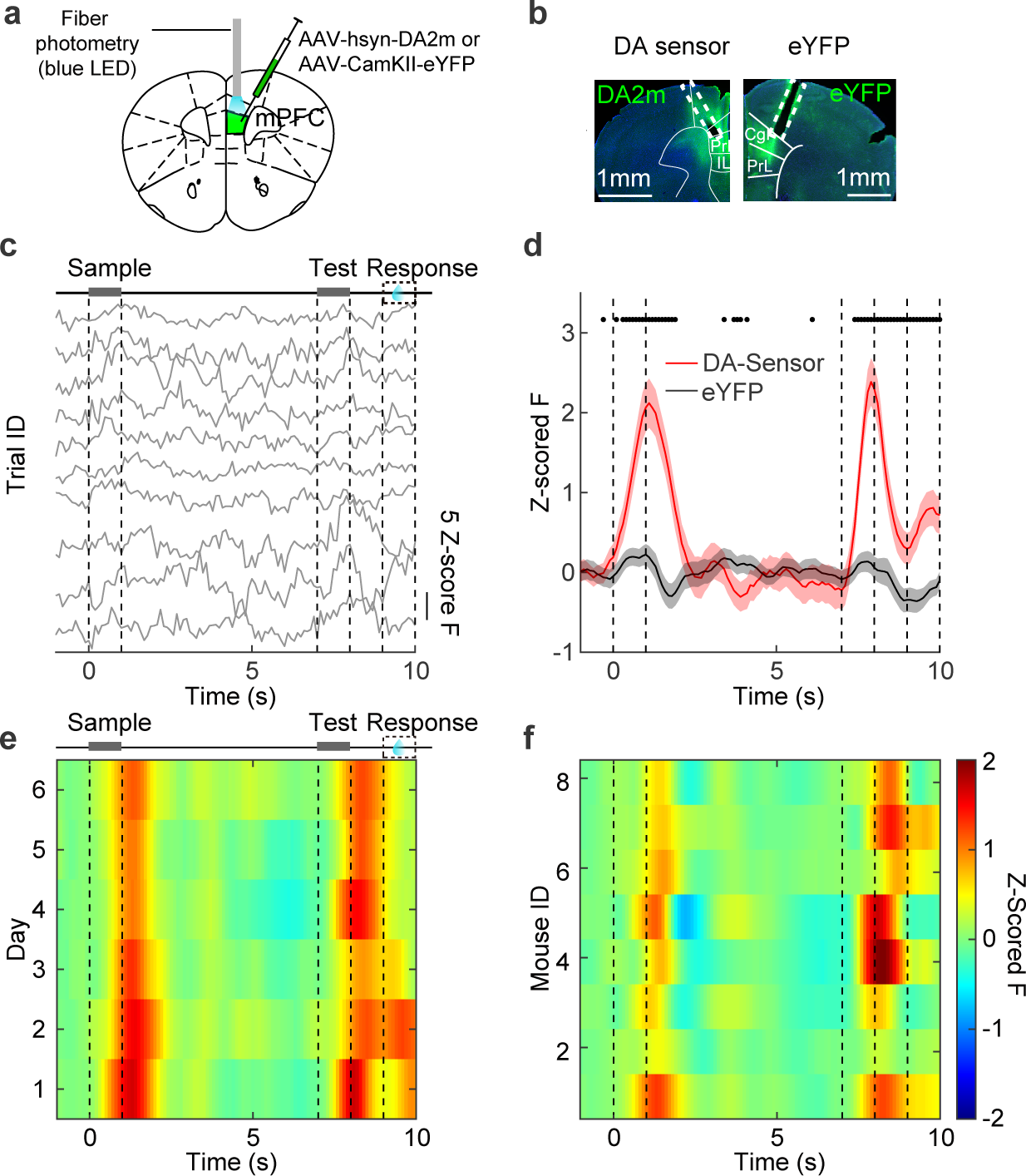
Transient dopamine dynamics in mPFC associated with WM. **a**, Strategy for fiber-photometry imaging of dopamine release in mPFC. **b**, Targeted expression of GRABDA sensor (DA2m) and eYFP and locations of optical fibers (white dashed boxes). Scale bar: 500μm for all figures. **c**, Z-scored GRABDA sensor signals of ten example trials from one recording session, with mean and standard deviation (STD) estimated from the baseline-period activity across trials. **d**, Averaged dopamine-sensor (red) and eYFP (black) signals of 200 trials for the recording session in **e**. **f**, Heatmap of averaged dopamine-sensor signals for the 6 training days of the mouse in **e**. **g**, Heatmap of averaged dopamine-sensor signal for 8 mice (averaged cross 6 sessions for each mouse).

Although individual traces fluctuated across trials (Fig.2c), on average sample-odor delivery elicited a transient increase in fluorescence during the early-delay period (Fig.2d). Such transient dopamine increase during the early-delay period was stable throughout training (Fig.2e, Extended Data Fig.7a-d) and across mice (Fig.2f). We observed significant differences in dopamine-fluorescence signals following the sample odors, especially early in learning (Extended Data Fig.7e), but no difference following the test odors (Extended Data Fig.7f-g). Consistent with its importance in reward and reward-prediction error^16,17^, dopamine responses were also larger in reward-delivered period than the non-rewarded condition (Extended Data Fig.7h). There was no difference in response latency for these conditions (Extended Data Fig.7i-l). These results indicate that endogenous dopamine activity in the mPFC is transiently modulated by WM-related stimuli, especially during the sensory-delivery and early-delay periods.

### Timing-dependent and bidirectional modulation of WM performance by dopamine in mPFC

Because dopamine elevation was predominantly increased during the sensory-delivery and early-delay periods (Fig.2), we tested whether behavioral performance could be differentially affected by elevating dopamine levels in the mPFC during different timing. Virus of AAV-DIO-ChR2 was bilaterally injected in the VTA of DAT-Cre mice to express ChR2 specifically in the dopaminergic neurons (Fig.3a-b). Optical fibers were bilaterally implanted in the mPFC of these mice to locally excite the dopaminergic terminals. Strikingly, WM performance was enhanced and impaired following laser illumination during the early- and late-delay period, respectively (Fig.3c). The behavioral changes were mainly due to the correct-rejection rate (Fig.3d). There was no change in the hit rate (Fig.3d) upon optogenetic excitation of dopaminergic terminals, therefore dopaminergic release in the mPFC did not modulate licking action *per se*.

**Fig.3.**
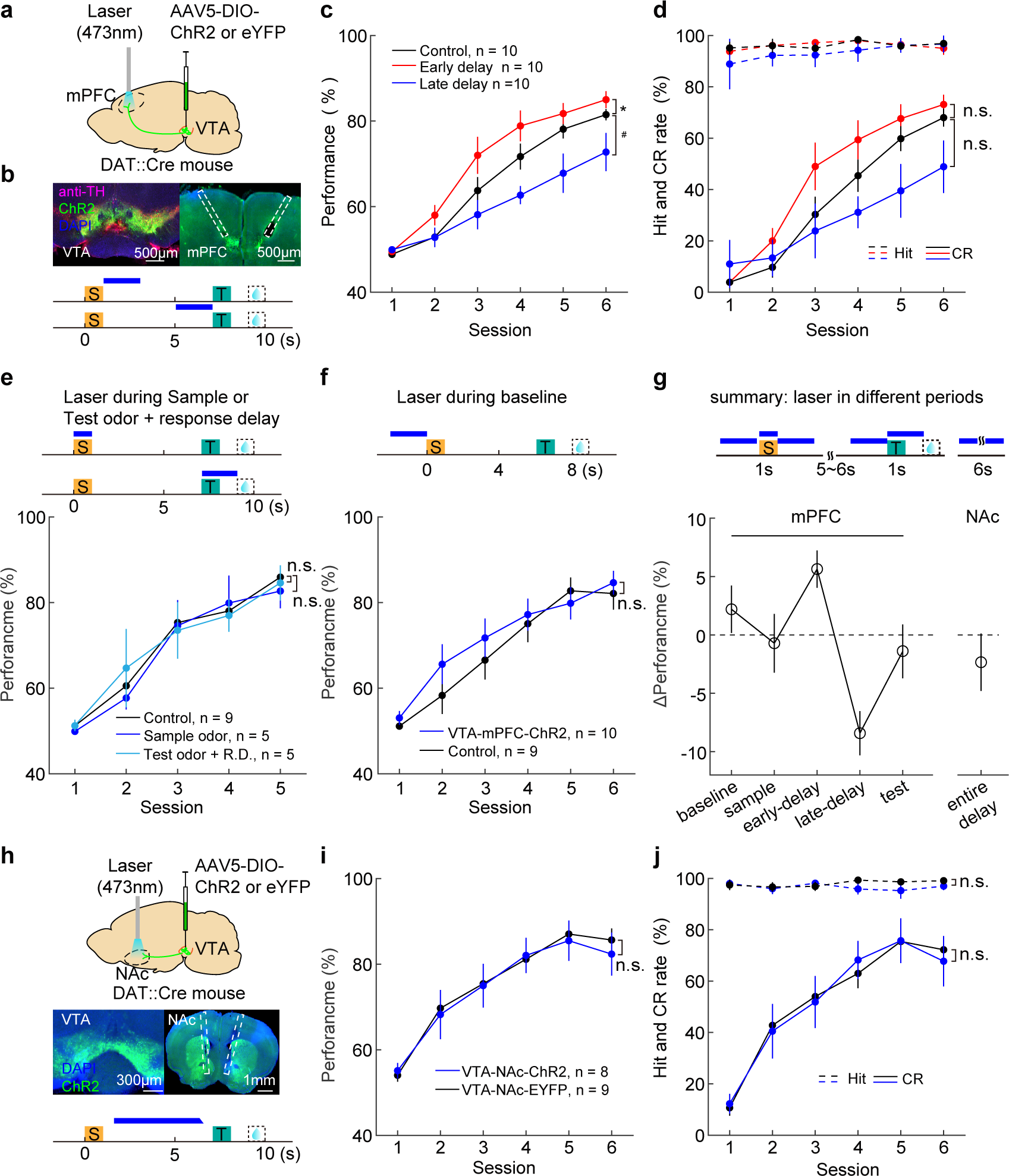
Timing-dependent and bidirectional modulation of WM by dopaminergic projections in mPFC. **a**, Strategy for optogenetic excitation of VTA dopaminergic terminals in mPFC. **b**, Above: Immunohistochemistry verification of virus expression in VTA (left) and optical fiber location (right). Below: Schematic diagram for optogenetic excitation of dopaminergic terminals in mPFC during the early- and late-delay period in ODPA task. **c**, Performance for the three groups of mice. For the control mice, one half with laser on during the early-delay period, and another half with laser on during the late-delay period. * p = 0.040; # p = 0.030. **d**, Hit and CR rates of mice as in **c**. **e**, Schematic diagram (above) and performance (below) for optogenetic excitation during either the sample- or test-odor period. For control group, laser was delivered during the sample-odor and test-odor/response-delay period for 4 and 5 mice, respectively. **f**, Schematic diagram (above) and performance (below) for optogenetic excitation during the baseline period. **g**, Summary of the temporal specificity in optogenetic excitation of dopaminergic projections. Performance of the training 4 days from 3^rd^ to 6^th^ was plotted. **h**, Above: Strategy for optogenetic excitation of VTA dopaminergic terminals in mPFC. Middle: Immunohistochemistry verification of virus expression in VTA (left) and optical fiber location (right). Below: Schematic diagram for optogenetic activating dopaminergic terminals in the NAc during the delay period in ODPA task. **i**, Performance for mice with optogenetic activation of dopaminergic terminals in NAc. **j**, Hit and CR rates of mice as in **i**.

We next examined the effect of constant elevation of dopamine during WM delay, by shedding blue light during the entire delay period, in DAT-Cre mice with AAV-DIO-ChR2 bilaterally injected in the VTA. We found a statistically insignificant trend of increase and decrease in performance for the task of 6-sec and 12-sec delay duration, respectively (Extended Data Fig.8a-b,d-i).

To examine whether suppression of dopamine release modulates behavioral performance, we injected the virus of AAV-DIO-NpHR3.0 in the mPFC (Extended Data Fig.8a,c). Optogenetic suppression of dopaminergic terminals by green laser during the delay period did not influence WM performance (Extended Data Fig.8j-l). Thus, dopaminergic projections to other regions might compensate for the loss of dopamine in the mPFC.

To further examine the temporal specificity of optogenetic manipulation, we excited the dopaminergic terminals in the mPFC during the sample-delivery, test-delivery, or baseline periods. No change in behavioral performance was observed under these conditions (Fig.3e-f). To test the spatial specificity of dopaminergic innervation for WM modulation, we also implanted optical fibers in the nucleus accumbens (NAc), which receives considerable dopaminergic innervation^46–48^. We did not observe any behavioral modulation following optogenetic excitation of dopaminergic terminals in the NAc (Fig.3h-j). The results of optogenetic manipulation (Day 3-6 in learning) were summarized in Fig.3g, showing the timing and spatial specificity in dopamine modulation of WM.

Dopamine modulates WM in a novel rather than familiar association^6^, so here we examined whether learning experience plays important role in dopamine modulation. After mice were over trained (the correct rate reached above 80% in one training session), activation of dopamine projections did not influence WM performance during either early- or late-delay period (Extended Data Fig. 8m-o). The learning-specific effect was consistent with the importance of dopamine in learning^21^, as well as our previous finding that suppressing the delay-period activity of the mPFC during the learning but not well-trained phase impaired WM behavioral performance^11^.

Taken together, these results demonstrated that dopamine modulation in WM behavior critically depends on the timing and location of dopamine release (Fig.1f-g,3g).

### Neural correlates of timing dependent-dopamine modulation in WM

To examine the neural correlates of timing-dependent dopamine modulation in WM, we performed simultaneous electrophysiological recordings and optogenetic excitation of dopaminergic terminals in the mPFC. Virus of AAV-DIO-ChR2 was injected in VTA of DAT-Cre mice and op-tetrodes^11,38^ were implanted in the mPFC, with electrodes inserted in both hemisphere of the mPFC and optical fiber inserted in the middle of two hemispheres (immunohistochemistry results from the mPFC and the VTA in Fig.4a, Extended Data Fig.9a). Mice were trained in the ODPA task and blue laser was illuminated during the delay period in a block design (20 trials per block, schematic diagram in Fig.4b), composed of three types of blocks: laser-off trials, early-excitation trials (laser on during early delay period), and late-excitation trials (laser on during the late delay period).

**Fig.4.**
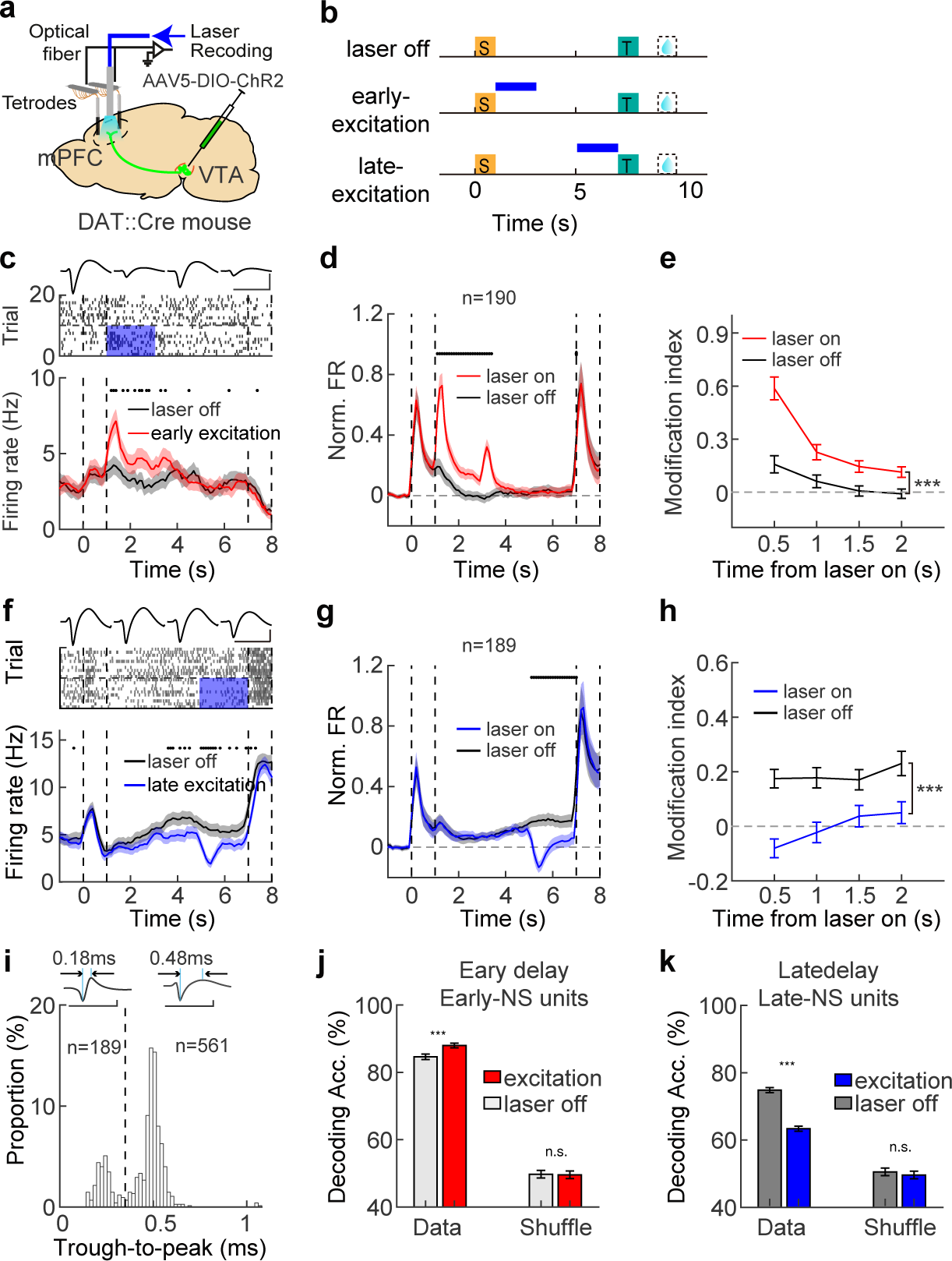
Modulation in mPFC neuronal activity by optogenetic excitation of dopaminergic projections. **a**, Strategy for optogenetic excitation of VTA dopaminergic terminals while simultaneously recording mPFC neuronal activity. **b**, Schematic diagram for three trial types: laser-off, early-excitation (laser-on in the early-delay period), and late-excitation (laser-on in late-delay period). **c**, Raster plot (above) and PSTH (below) showing activity of an example neuron with increased firing rate (FR) by optogenetic excitation during the early-delay period. Top curves show averaged spiking waveforms recorded from tetrodes. Blue shades indicate laser illumination. **d,** Averaged normalized firing rate (Norm. FR) for early-increased units (n = 190). **e**, Modulation index within 500-ms bin during the early-delay period for the early-increased units in the laser-off and laser-on trials in **d**. ***p < 0.001, Wilcoxon rank sum test. **f**, As in **c** for optogenetic excitation during the late-delay period. An example neuron showing decreased firing rate during laser illumination. **g**, Averaged normalized firing rate for late-decreased (below, n = 189) units. **h**, Modulation index within 500ms bin in late delay period for the late-decreased units in the laser-off and laser-on trials in **g. i,** Distribution for trough-to-peak of all recorded units (n = 750). Dashed line indicates 0.35ms. Above: Waveforms for two example neurons with trough-to-peak of 0.18 ms and 0.48 ms. NS: narrow-spiking neurons (n = 189). BS: broad-spiking neurons (n = 561). **j**, SVM decoding accuracy for NS units from early-changed (Early-NS) units during the early-delay period (n = 92). **k**, SVM decoding accuracy for NS units from late-changed (Late-NS) units during the late-delay period (n = 112).

We observed significant modulation in neuronal firing by optogenetic excitation. The activity of mPFC neurons could be either elevated (Fig.4c) or reduced (Extended Data Fig. 9e) by early-excitation of dopaminergic terminals (heatmap in Extended data Fig.9b). Overall, 25% (190 out of 750) and 22% (165 out of 750) of the mPFC neurons exhibited increased and decreased firing modulation upon early excitation of dopaminergic terminals, respectively (termed ‘early-increased’ and ‘early-decreased’, Extended data Fig.9d). Notably, the early-increased neurons showed higher delay activity with optogenetic excitation than that without laser illumination in control trials (Fig.4d). Optogenetic excitation of dopaminergic terminals during the early-delay period enhanced the gain of endogenous delay activity for these neurons (Fig.4e). For other neurons with reduced activity following early-delay optogenetics, we did not observe significant delay-period activity (Extended data Fig.9f-g).

We then examined mPFC neural correlates following optogenetic dopaminergic excitation during the late-delay period. As shown in Fig.4f Extended data Fig.9i, the activities of the example neurons were elevated or reduced by late-excitation (heatmap in Extended data Fig.9c). Overall, 32% (243 out of 750) and 25% (189 out of 750) of the mPFC neurons exhibited increased and decreased firing modulation upon late delay-optogenetic excitation of dopaminergic terminals, respectively (termed ‘late-increased’ and ‘late-decreased’, Extended data Fig.9h, heatmap in Extended data Fig.9h). Importantly, the late-decreased neurons exhibited elevated activity comparing to baseline (black curve, Fig.4g). Thus, late-delay optogenetic dopaminergic excitation reduced the endogenous delay-period activity that was associated with WM (Fig.4h). Conversely, in the late-increased neurons, we observed no modulation in endogenous delay-period activity comparing to baseline (Extended data Fig.9j). Thus, these late-increased neurons were irrelevant to WM endogenously during the late-delay period, and the increase in their activity could act as noise (Extended data Fig.9k).

Furthermore, our observations revealed that dopamine differentially influenced the activity of distinct mPFC neurons during various phases of the delay period. Specifically, certain neurons exhibited modulation solely during the early-delay excitation, as depicted in Extended Data Fig.10g (left). Conversely, a separate set of neurons displayed modulation exclusively during the late-delay excitation, illustrated in Extended Data Fig.10g (middle). There were also neurons that underwent modulation from both early- and late-delay excitations, as depicted in Extended Data Fig.10g (right). The proportions of these distinct neuron groups are shown in Extended Data Fig.10h-i. These results demonstrated that dopamine modulates neuronal activity in a context-dependent manner.

We further tested the potential changes to WM information-encoding ability of mPFC neurons following optogenetic early- and late-excitation of dopamine projections. Indeed, mPFC neurons could change their coding ability (Extended data Fig.10a,d), in a manner dependent on cell types. Based on trough-to-peak distribution, we clustered cells into broad spiking (BS, putative excitatory neurons, 74.8%) and narrow spiking (NS, putative inhibitory neurons, 25.2%; Fig.4i). We focused on the BS and NS neurons with significant modulation by laser illumination. For the BS neurons, both early- and late-delay excitation slightly improved coding ability (Extended data Fig.10b-f). For the NS neurons, early- and late-delay excitation respectively improved and impaired WM-coding ability (Fig.4j-k), in manner consistent with behavioral modulation (Fig.3).

In summary, early- and late-delay excitation of dopaminergic projection from the VTA to the mPFC increased and decreased the gain of WM relevant activity, respectively. Late-delay excitation also evoked more noise. Dopamine excitation during the early- and late-delay period also bi-directionally modulated coding ability for WM information in the putative inhibitory neurons, in a manner consistent with modulation in behavioral performance.

## Discussion

Many previous studies have demonstrated the importance of dopamine neurons in WM and its related delay-period activity. However, the timing-specific requirement of dopamine action on WM and its neural mechanisms remain unclear. We tackled these questions in mice performing an olfactory WM task by combing dopamine sensor imaging, pathway-specific optogenetics and electrophysiological recordings. We found that VTA dopamine neurons critically modulate WM performance and could transiently encode WM information (Fig.1). Dopamine signals in the mPFC were only transiently elevated during the early-delay period in performing the WM task (Fig.2). Dopamine release in mPFC modulates WM behavioral performance in a timing-dependent manner, as shown by improved and impaired performance following optogenetic dopaminergic excitation during the early- and late-delay period, respectively (Fig.3). Such bidirectional modulation in performance was associated with respective increase and decrease in activity gain and coding ability of some mPFC neurons (Fig.4).

Previous studies using pharmacology methods have demonstrated that the neuromodulation of dopamine in modulating WM performance and its related neuronal activities depend on the concentration of dopamine an inverted-U shape^40–43^, which we also validated by optogenetic manipulation of delay-period activity of dopaminergic neurons (Fig.1f-g). In the current study, we further discovered the importance of release timing in dopamine modulation of WM performance. We found that the endogenous pattern of dopamine release in mPFC was transient increase during the early-delay period, without significant changes during the late-delay period (Fig.2). Importantly, only dopamine elevation confirming the endogenous patterns could improve WM performance (Fig.3), stressing the importance of observing neuromodulator dynamics at a high temporal resolution. Such timing-specific modulation by dopamine suggests potential new methods in deep-brain stimulation to rescue WM deficits accompanying many brain disorders, such as schizophrenia^49^ and Parkinson’s disease^50^.

A hallmark of WM is resistance to distractors during the delay period^49^. Our previous report show that distractor application during the delay period could elicit transient activity of insular cortical neurons to resist distractor^39^. A salient distractor might capture attention of subjects and induce neuromodulator release. Such distractor induced neuromodulation might be related to our finding that the late-delay excitation of dopaminergic terminals impaired WM performance (Fig.3). Preventing such increase in dopamine during the late-delay period might serve for robust information maintenance upon distractor.

Brain-slice recordings have revealed diverse effect of ion channels and synaptic changes upon dopamine treatment^51–54^. Physiological studies *in vivo* have shown that dopamine transients in mPFC could increase firing^18,20^ or increase activity gain to help associative learning_16,55_. It remains to be determined whether and how the abovementioned ion-channel and synaptic mechanisms contribute to our observed cell-type and delay-phase dependent modulation in WM performance and its associated activity.

Dopamine exerts diverse functions in the brain, for example reward prediction error and motivation^14–17^ and motor action^22^. Our optogenetic excitation of the VTA-to-mPFC projections during the decision-making and reward periods did not significantly modulate WM behavioral performance (Fig.3e). Therefore, modulation of behavioral performance during the delay-period excitation is unlikely to be attributed to factors like reward prediction error, motivation, or motor actions. It is plausible that dopaminergic projections targeting other brain regions might be responsible for facilitating the functions mentioned above within their respective phases.

Dopaminergic inputs to the PFC innervate both excitatory and inhibitory neurons^56^. Moreover, pharmacological perturbation of dopamine receptors in PFC differentially modulates coding ability of WM information in putative excitatory and inhibitory neurons^57^. Optogenetic perturbation show that dopamine from the VTA modulates activity of inhibitory neurons in the mPFC to mediate conditional learning^58^. Here we found that coding ability of putative inhibitory neurons was modified by early- and late-delay optogenetics in a manner consistent with behavioral modulation (Fig.4i-k). Future studies are required to dissect the potential role of different types of inhibitory neurons in dopamine modulation of behavior.

In conclusion, dopamine in mPFC modulates working memory in a timing-dependent manner, accompanied by modification of activity gain and memory-coding ability in mPFC neurons. Our findings underscore the importance of timing in dopamine modulation of the brain.

**Extended Data Fig.1| Behavioral performance during learning. a**, Behavioral performance for ODPA task in 6 consecutive learning days. Error bars indicate SEM unless stated otherwise. **b**, Lick raster of 50 trials of an example session for an example mouse. Each tick represents for one lick. Blue, black, and red dots on the right indicate for the correct rejection, hit, and false alarm trials, respectively. **c**, Immunohistochemistry verification of virus (NpHR or ChR2) expression and optical fiber locations in VTA. **d**, Hit and CR rates with optogenetic suppression during the delay period. **e**, Hit and CR rate with optogenetic excitation during the delay period. **p < 0.01, Two-way ANOVA with repeated measure unless stated otherwise. **f**, Hit and CR rates with optogenetic suppression during the baseline period. **g-i**, Lick rates of mice as in **d-f** in the 3^rd^ learning day. Black dots above the curves indicate p < 0.05 within 100-ms bin. Wilcoxon rank sum test unless stated otherwise.

**Extended Data Fig.2| Behavioral results for optogenetic excitation of VTA^DA^ and VTA^GABA^ neurons in learning the ODPA task. a**, Performance for optogenetic excitation of VTA^DA^ neurons during the intertrial interval. **b**, Hit and CR rate for the mice in **a**. **c**, Lick rates of the mice in **a** in the 3^rd^ learning day. **d** and **g**, Performance for optogenetic excitation of VTA^DA^ neurons during the delay period at a higher laser power than in Figure **1c** (2 mW and 4 mW vs. 1 mW). **e** and **h**, Hit and CR rate for mice in **d** and **g** (*p = 0.026 and 0.024 for Hit and CR with 2mW laser; *p = 0.0018 and 0.012 for Hit and CR with 4mW laser). **f** and **i**, Lick rates of mice in **d** and **g** in the 3^rd^ learning day. **j**, Performance for optogenetic excitation of VTA^GABA^ neurons during the delay period. **k**, Hit and CR rate for mice in **j**. **l**, Lick rates of mice in **j** in the 3^rd^ learning day.

**Extended Data Fig. 3| Population calcium signals of VTA^DA^ neurons in OPDA task. a**, Strategy for monitoring the activity of VTA^DA^ neurons using fiber photometry (top) and immunohistochemistry verification of virus (GCaMP6s) expression and optical fiber locations in VTA (bottom). **b**, Example traces for calcium signals of VTA^DA^ neurons in ODPA task. Each trace indicates one trial. **c**, PSTH for averaged calcium signals of VTA^DA^ neurons. **d**, Calcium signals of VTA^DA^ neurons for different sample odor (sample odor1: S1; sample odor2: S2). **e-f**, Heatmap showing calcium signals of VTA^DA^ neurons in S1 (**e**) and S2 (**f**) trials cross 6 training sessions for 14 mice. Each line indicates the results from one mouse. **g**, Calcium signals of VTA^DA^ neurons for the reward and no-reward trials. **h-i**, Heatmap showing calcium signals of VTA^DA^ neurons in the reward (**h**) and no-reward (**i**) trials cross 6 training sessions for 14 mice. **j-k**, Performance (**j**) and Hit and CR rate (**k**) for mice with fiber photometry imaging in **a-i**. **l**, Peak response for calcium signals of VTA^DA^ neurons in the rewarded and no-rewarded trials, two seconds following the start of response window, for 6 training sessions. ***p < 0.001. **m**-**o**, Peak response for calcium signals of VTA^DA^ neurons to different sample odors (**m**, within two second after onset of sample delivery), test odors (**n**, within two seconds after onset of test-odor delivery) and paired odors (**o**, within two seconds after onset of test-odor delivery; paired trials, S1-T1 or S2-T2; non-paired trials, S1-T2 or S2-T1) in 6 training sessions.

**Extended Data Fig.4| Electrophysiological recoding in VTA. a**, Immunohistochemistry verification of virus (ChR2) expression and optrodes locations in VTA. **b**, PSTHs for two example neurons in VTA showing the firing rate to sample odors. **c**, PSTH for two example neurons showing the firing rate in Hit/FA and Hit/CR trials. **d**, Heatmap showing selectivity for the selective VTA^DA^ neurons (n = 13, out of 23).**e**, Heatmap showing activity modulation for the selective neurons in VTA (n = 332, out of 1251). **f**, Decoding accuracy of the neurons in **e**. Decoding method: maximum correlation coefficient (MCC) with 500 repeats.

**Extended Data Fig. 5| Electrophysiological recoding in mPFC with optogenetic excitation of VTA^DA^ neurons during the delay period. a**, Above: Immunohistochemistry verification of optrodes in mPFC (left), as well as ChR2-virus expression and fiber locations in VTA (right). Below: Diagram of laser on/off block design for optogenetic excitation of VTA^DA^ neurons during the delay period. **b-c**, Raster (above) and PSTH (below) showing activity of two example mPFC neurons with (blue) and without (black) optogenetic activation of VTA^DA^ neuron. **d-g**, PSTH showing activity of four extra example mPFC neurons for the laser-on (blue) and laser-off (black) blocks, showing prominent modulation for these neurons, even during the baseline period. **h-o**, PSTH showing activity of eight extra example mPFC neurons, showing modulation in coding ability and firing rate following optogenetic excitation of VTA^DA^ neurons during the delay period. Above: laser off. Below: laser on. **p**, Activity heatmaps showing the mPFC units with reduced activity in the laser-on (right) trials comparing to the laser-off (left) trials (n = 293). **q**, PSTH of normalized firing for the neurons in **p**. **r**, SVM decoding accuracy of mPFC neurons with reduced firing following activation of VTA^DA^ neurons. **s-u**, as in **p-r** for all the recorded mPFC neurons (n = 475).

**Extended Data Fig. 6| Electrophysiological recoding in mPFC with optogenetic suppression of VTA^DA^ neurons during the delay period. a-u**, as in Extended Data Figure 5, for expression of NpHR in VTA^DA^ neurons.

**Extended Data Fig.7| Behavioral performance and dopamine response in mPFC during learning. a**, Behavioral performance for mice in fiber photometry-imaging experiments. **b**, Hit and correct rejection rates for mice in **a**. **c**, Licking raster for an example mouse during imaging. Each tick represents for one lick. Black, blue, and red dots on the right indicate for the hit, correct rejection, and false alarm trials, respectively. **d**, Averaged dopamine-sensor signals for 8 mice in 6 recording sessions. **e**, Schematic diagram of ODPA task (up) and the peak of GRABDA response to sample odors for 8 mice in 6 training sessions (below). The response peaks were calculated in 2-s duration following the sample onsets as shown in the red dotted box. *p = 0.036, Two-way ANOVA with repeated measure test was performed cross sessions between stimuli. **f-h**, As in **e** for test odors, reward prediction odors and reward. *p = 0.019 in **h**. **i**-**l**, Latency of the response peaks from stimuli onset in **e**-**h**.

**Extended Data Fig.8| Behavioral performance of optogenetic manipulation on VTA dopaminergic terminals in mPFC. a**, Strategy for optogenetic excitation or suppression of VTA dopaminergic terminals in mPFC. **b**, Immunohistochemistry verification of virus (ChR2) expression in VTA and optical fiber locations in mPFC. **c**, Histochemistry verification of virus (NpHR) expression in VTA and optical fiber locations in mPFC. **d**, Schematic diagram (top) and performance (below) for optogenetic excitation during 6-s delay period. **e**, Hit and CR rates of mice as in **d**. *p = 0.030 for CR. **f**, Lick rates of mice as in **d** and **e** (in #3 training session). **g**, Schematic diagram (top) and performance (below) for optogenetic excitation during 12-s delay period (p = 0.093). **h**, Hit and CR rates of mice as in **g** (*p = 0.021 for CR). **i**, Licking rates of mice as in **g** and **h** (in #3 training session). **j,** Schematic diagram (top) and performance (below) for optogenetic inhibition during 5-s delay period. **k**, Hit and CR rates of mice as in **j**. **l**, Licking rates of mice as in **j** and **k** (in #3 training session). **m**, Schematic diagram (top) and performance (below) for optogenetic excitation during early and late delay period. For the control mice, laser was delivered during early-delay period for one half and late-delay period for the other half. **n**, Hit and CR rates of mice as in **m**. **o**, Licking rates of mice as in **m** (in 3rd training session).

**Extended Data Fig. 9| Neural modulation of mPFC neurons induced by optogenetic excitation of dopaminergic terminals. a**, Immunohistochemistry verification of virus (ChR2) expression in VTA and optrode locations in mPFC**. b**, Heatmaps showing neural modulation by early-delay optogenetics, for the neurons with increased (above, n = 190) and decreased (below, n = 165) firing. Left, laser-off trials; right, early-excitation trials. **c**, as **b** for modulation by late-delay optogenetics. Firing-increased neurons: above, n = 243. Firing-decreased neurons: below, n = 189. **d**, Proportion of neuronal modulation following early-excitation of dopaminergic terminals in mPFC. **e**, Raster plot (above) and PSTH (below) showing activity of an example neuron with decreased firing rate by optogenetic excitation during the early-delay period. Top curves show averaged spiking waveforms recorded from tetrodes. Blue shades indicate laser illumination. **f,** Average normalized firing rate (Norm. FR) for the early-decreased (n = 165) units. **g**, Modulation index within 500-ms bin during the early-delay period for neurons in **d**. ***p < 0.001, Wilcoxon rank sum test. **h-k**, as in **e-g** for late-excitation of dopaminergic terminals in mPFC.

**Extended data Fig.10| Modulation in memory-coding ability of mPFC neurons by optogenetic excitation of dopaminergic terminals. a**, PSTH showing activity of three example neurons with increased (left and middle) and decreased (right) coding ability for sample odors with optogenetic excitation during the early-delay period (early excitation). Above: laser-off trials. Below: early-excitation trials. Blue shadows: laser delivery. **b**, Waveform trough-to-peak distribution for the neurons with significant firing modulation following early-excitation. **f**, Decoding accuracy for early-changed broad spiking (BS) neurons (n = 263). **d-f**, as in a-d for late-excitation. Narrow spiking (BS) neuron number: n = 320. **g**, PSHT showing activity of six example neurons with significant firing modulation by laser-excitation. Left two neurons show firing increased (above) or decreased (below) only in early-excitation trials. Middle two neurons show firing increased (above) or decreased (below) only in late-excitation trials. Right two neurons show firing increased (above) or decreased (below) in both early- and late-excitation trials. **h**, Proportion for firing increased or decreased units in each 500-ms bin during 2-s laser illumination period. **i**, Proportion of overlap for firing changed units during the early- or late-excitation condition.

## MATERIALS AVAILABILITY

Further information and requests for resources and reagents should be directed to and will be fulfilled by the Lead Contact, Chengyu T. Li. The study did not generate new unique reagents.

## Methods

### Animals

Dopamine transporter (DAT)::IRES-Cre knockin mice^59^ (referred to as DAT-Cre mice hereafter) were obtained from the Jackson Laboratory (JAX006660, RRID:IMSR_JAX:006660) and crossed with WT C57/BLJ6 mice. Male DAT-Cre mice (mostly heterozygous) were used for optogenetic experiments and their DAT^-/-^ littermates were used as control. Male DAT-Cre mice were used for fiber photometry and electrophysiological recording experiments. Surgeries were performed for mice between 8-12 weeks of age and between 25 to 35g in weight. Behavioral training started a month after surgery. All mice were housed with *ad libitum* access to water and food at a stable 12-h light-dark cycle (light on from 7:00 a.m. to 7:00 p.m.). All animal experimental procedures in current study were approved by the Animal Care and Use Committee of the CAS Center for Excellence in Brain Science and Intelligence Technology, Shanghai, China (Table 1).

**Table 1.**
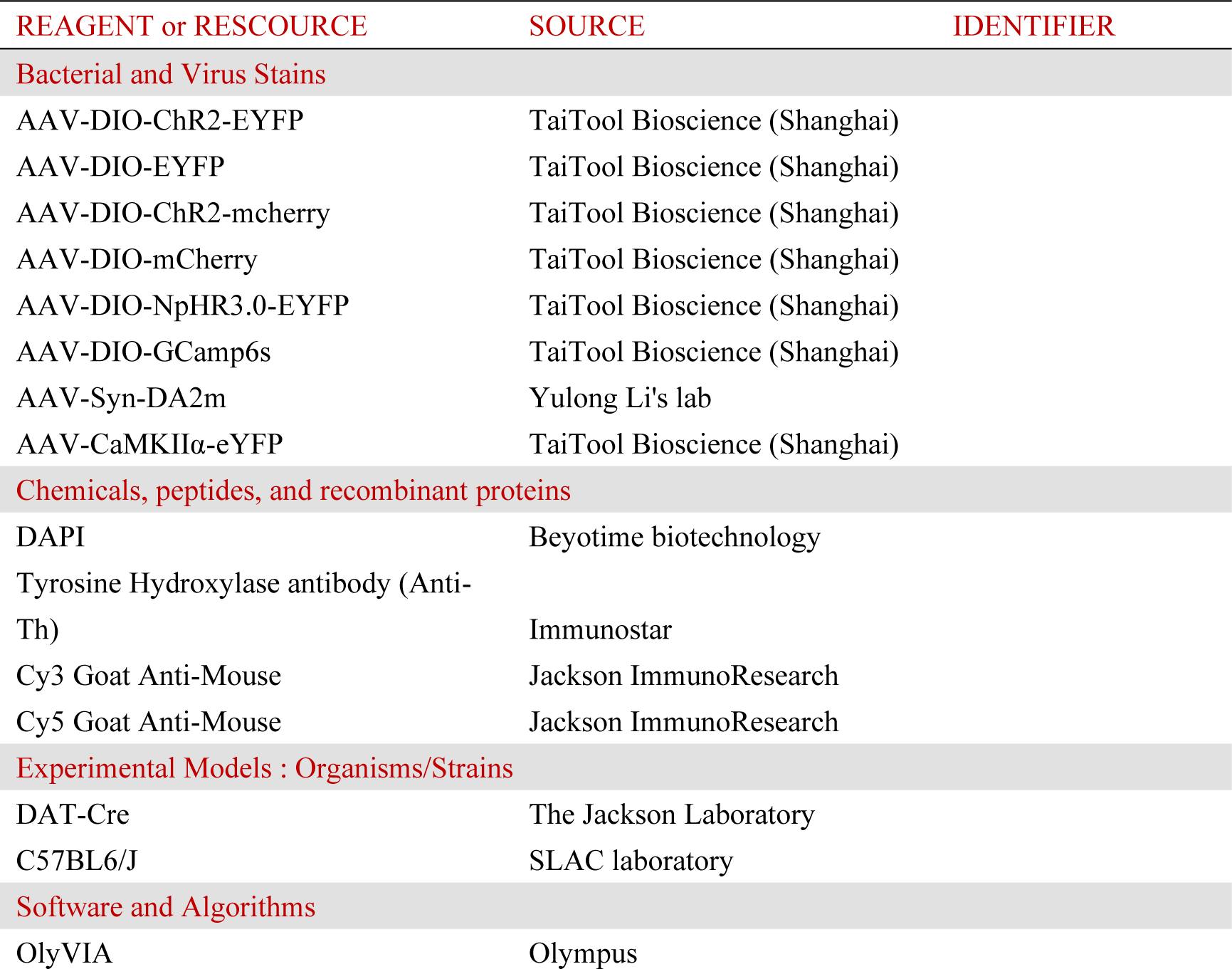
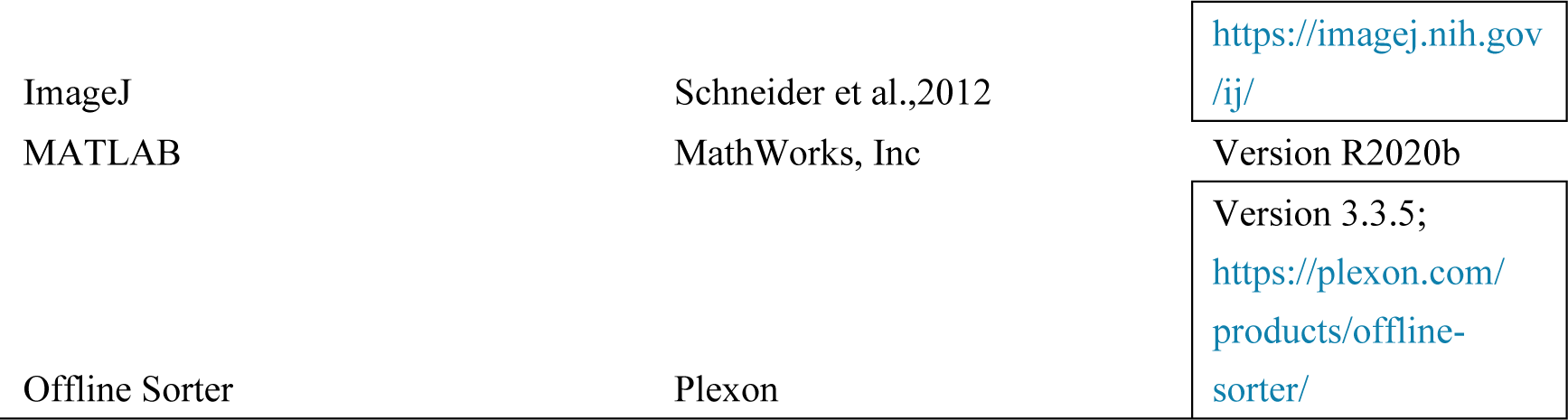

| REAGENT or RESOURCE | SOURCE | IDENTIFIER |
| --- | --- | --- |
| **Bacterial and Virus Stains** |  |  |
| AAV-DIO-ChR2-EYFP | TaiTool Bioscience (Shanghai) |  |
| AAV-DIO-EYFP | TaiTool Bioscience (Shanghai) |  |
| AAV-DIO-ChR2-mcherry | TaiTool Bioscience (Shanghai) |  |
| AAV-DIO-mCherry | TaiTool Bioscience (Shanghai) |  |
| AAV-DIO-NpHR3.0-EYFP | TaiTool Bioscience (Shanghai) |  |
| AAV-DIO-GCamp6s | TaiTool Bioscience (Shanghai) |  |
| AAV-Syn-DA2m | Yulong Li's lab |  |
| AAV-CaMKIIα-eYFP | TaiTool Bioscience (Shanghai) | |
| **Chemicals, peptides, and recombinant proteins** |  |  |
| DAPI | Beyotime biotechnology |  |
| Tyrosine Hydroxylase antibody (Anti-Th) | Immunostar |  |
| Cy3 Goat Anti-Mouse | Jackson ImmunoResearch |  |
| Cy5 Goat Anti-Mouse | Jackson ImmunoResearch |  |
| **Experimental Models : Organisms/Strains** |  |  |
| DAT-Cre | The Jackson Laboratory |  |
| C57BL6/J | SLAC laboratory |  |
| **Software and Algorithms** |  |  |
| OlyVIA | Olympus |  |

### AAV construction

The vectors of optogenentic experiments were obtained from AddGene. Recombinant adeno-associated virus serotype 5 (AAV2/5) vectors containing ChR2 and NpHR (AAV2/5-hEF1a-DIO-hChR2(H134R)-EYFP-WPRE-pA, AAV2/5-EF1a-double floxed-hChR2(H134R)-EYFP-WPRE-HGHpA, AAV2/5-hEF1a-DIO-eNpHR 3.0-EYFP-WPRE-pA) or fluorescent proteins (AAV2/5-EF1a-double floxed-EYFP-WPRE-HGHpA, AAV-hsyn-eGFP-3Flag-WPRE-SV40pA) were packaged by Shanghai Taitool Bioscience Co. Ltd and Gene Editing Core Facility of CEBSIT, Institute of Neuroscience, CAS, Shanghai. The virus AAV2/9-ayn-Flex-GCaMP6s-WPRE-pA and AAV2/9-Syn-DA2m^12^ for fiber photometry experiments were obtained from Shanghai Taitool Bioscience Co. Ltd and Yulong Li’s lab of Peking University, respectively. The source of all virus was listed in Table 1.

### Surgery

All surgeries were performed under anaesthetized and aseptic conditions. Body temperature was maintained with a heating pad below and sterile cloth covering mouse. Before surgery, mice were anaesthetized in a box added isoflurane (0.6 mL/L) for induction in fume cupboard. Within 2 minute, anesthetic mice were moved to a digital small-animal stereotaxic instrument (Shenzhen Ruiwode Lift Technology Co.,Ltd.) and kept anesthesia with isoflurane mixed with oxygen (0.8 ∼ 1.0 %, 0.4 ∼ 0.6 L min oxygen flow rate). Head skin was sterilized with iodophor. Head hair was removed from dorsal surface after initial induction with curved ophthalmic scissors. Eyes were protected with ophthalmic ointment during surgery. The head skin was removed and the tissue on the skull was cleaned up with sterile swabs. Head skull was adjusted to horizontal according to bregma and lambda levels, and location of targeted brain areas were marked on skull. Then the plates to fixed mice head were glued on skull with tissue glue and fastened with dental cement from the edge without affecting target markers. The craniotomy (0.5-1 mm in diameter) for virus injection above targeted areas was performed by cranial drill, submerged in artificial cerebrospinal fluid without glucose. Virus injection was performed without removing dura and implemented by microinjection needles, of which the inner diameter of tip was 30 μm and the mouth wad 60° beveled. Needle was controlled by a 10 μl microsyringe pump (Shenzhen Ruiwode Lift Technology Co.,Ltd.) and speed of virus delivering was at a rate of 50 nl / min. Needle was kept still in brain for 10 minutes after completion of injection then was slowly withdrawn. Silicone elastomer (Kwik-Sil^TM^, WPI) was used for separating the incision area and dental cement to protect the exposed skull and dura. For implanting optical fibers or electrodes, the diameter of craniotomy was larger according to the inserted fibers or electrodes, and dura above brain tissue was peeled off. For injection or implantation at multiple locations, craniotomy of multiple sites was performed within a single surgery. Mice were intraperitoneally injected antibiotic drug after surgery for 3 days (Ampicillin sodium, 20 mg / ml, 160 mg / kg b.w.) and maintained on a heating pad for serval hours before moved into home cages. Histological verification was performed to confirm targets for virus injection and fiber implantation.

#### Optogenetic experiments

ChR2 or NpHR (titer: 3∼7 × 10^12^, 0.4 μl per site) was bilaterally injected in the VTA (bregma, AP: -3.30∼3.40mm, ML: ±1.30mm, DV: -4.30mm, angle: 10° to the lateral) of DAT-Cre mice and their control littermates. Optical fibers (200µm in diameter, 0.37 NA) were bilaterally implanted just 0.15mm above the injection sites for manipulating VTA^DA^ neurons or in mPFC (bregma, AP: +1.97mm, ML: ±1.26mm, DV: -1.70mm, angle: 26° to the lateral) or NAc shell (bregma, AP:+1.5mm, ML: ±1.35mm, DV: -4.3mm, angle: 8° to the lateral) for manipulating dopaminergic terminals. For control group mice, if the number of control littermates were not enough, DAT-Cre positive mice were injected control virus (AAV2/5-EF1a-double floxed-EYFP-WPRE-HGHpA, titer: 2∼5 × 10^12^) for balance.

#### Fiber photometry experiments

GCamp6s (titer: 3∼5 × 10^12^,0.4 μl) was unilaterally injected into VTA (bregma, AP: -3.30∼3.40mm, ML: +0.5mm, DV: -4.2mm) of DAT-Cre mice and an optical fiber was implanted just above the virus for monitoring population activity of VTA^DA^ neurons. For monitoring dopamine dynamics, dopamine sensor (AAV-syn-DA2m, titer: 3∼5 × 10^12^,0.4 μl) was injected into mPFC of DAT-Cre mice (AP: +1.97, ML: ±1.26, DV: -1.75, mm, angle: 26°) and optical fibers were implanted just above the virus. For GFP control, AAV-CamKII-GFP was injected into VTA or mPFC of C57 mice.

#### Electrophysiological recoding

AAV2/5-hSyn-DIO-ChR2-eYFP or and AAV2/5-hSyn-DIO-NpHR-eYFP was injected into the VTA of DAT-Cre mice. For recoding in VTA, homemade optrodes were inserted into VTA unilaterally. For recording in mPFC and optogenetic excitation of VTA^DA^ neurons or dopaminergic terminals in mPFC, electrodes were bilaterally inserted into mpFC and optical fibers were bilaterally injected into VTA or only optrodes were inserted in mPFC. For recording in mPFC and optogenetic inhibition of VTA^DA^ neurons, NpHR was unilaterally injected into VTA to ensure that mice could perform the task.

### Behavioral setups

Some of the following methods are like that in the previously published^37,38,60^. We utilized olfactory delayed paired association (DPA) task in head-fixed mice. Computer controlled olfactometry systems were used for automatic training. Odor delivery lasts for 1s and water reward is ∼4uL/drop. Licks were detected using capacitance module for optogenetic experiments and fiber photometry and using infrared blocking in electrical recording.

### Behavioral training

We utilized the DPA task for all optogenetic experiments and recording experiments, which was carried out as descried previously_37,38,60_. In brief, two different olfactory stimuli were delivered successively with a 5 - 12 sec delay period. We defined the first odor as sample and the second odor as test odor (odor list see table 2), each applied for 1 sec in duration. One sec after the offset of test odor, response window (1 sec in duration) started. Mice were trained to lick for water reward in paired trials (S1-T1, S2-T2) and not lick in non-paired trials (S1-T2, S2-T1). In the paired trials, licks in response window triggered ∼4 mL water delivery, which was marked as hit (Hit). No lick during response window resulted in reward omission and was marked as miss (Miss). In the non-paired trials, no water reward was delivered; licks and no lick were marked as false alarm (FA) and correct rejection (CR), respectively. Mice preformed 200 trials per day.

**Table 2.**

| Odor pairs |  |  |  |
| --- | --- | --- | --- |
| Sample 1 | Sample 2 | Test 1 | Test 2 |
| 1-Pentanol | Butyl formate | Ethyl acetate | 3-Methyl-2-buten-1-ol |
| Ethyl propionate | Pentyl acetate | Methyl butyrate | Propyl formate |
| 1-Pentanol | Butyl formate | Methyl butyrate | Propyl formate |

Mice were under water restriction through the whole training period and only got water in the training box. Mice got ∼20 drops water to habituate training setups at the beginning. Free water was available after the training in each day. Body weight was maintained above 80% of their full body weight. There were five phases for behavioral training: preparation, habituation, shaping, learning and well-trained. About 4 weeks of recovery from surgery, mice were water-deprived for 2 days in their home cages. Then mice were head fixed in training tubes to learn licking for water in the training box about 30minites. This habituation of the training system typically lasted for 3 days. After that the shaping process started, in which mice learnt to associate odor stimuli with water reward. Only the paired odors were presented, therefore, all trials could result in water reward. To realize automatically training, there were 10-40 teaching trials for the shaping period, in which water reward came out of water-port tip automatically in the response window no matter whether mice lick or not. Normally, it took 3 days for mice learned it well to lick water in response window after the second odor. Then, it entered learning phase, in which both paired and non-paired trials were presented. Mice learned to lick in the paired trials and not lick in nonpaired trials in ∼80% trials after 6 learning days. After that mice keep stable behavior above 80% in well-trained phase. In this study, we focus on the learning phase.

### Behavioral performance measurements

Correct rate = (num. hit trials + num. correct rejection trials) / total number of trials;

Hit rate = num. hit trials / (num. hit trials + num. miss trials);

Miss rate = num. miss trials / (num. hit trials + num. miss trials);

FA rate = num. false alarm trials / (num. false alarm trials + num. correct rejection trials);

CR rate = num. correct rejection trials / (num. false alarm trials + num. correct rejection trials);

Δ Performance = correct rate of group 1 – correct rate of group 2.

### Behavioral optogenetic experiments

#### Optogenetic excitation of VTA^DA^ neurons

To test the effect of activating VTA^DA^ neurons during different epoch of task, blue laser (473nm) was delivered during delay, sample, test, baseline and inter-trial-interval (ITI) period when mice performing the ODPA task. For excitation during delay, we activated VTA^DA^ neurons with laser illumination of 1, 2 and 4mW in a 5-second interval in the middle of 6 seconds. For excitation during sample odors, 1mw laser was delivered from the onset to the offset of sample odors. For excitation during test odors, 1 mw laser was delivered from the onset to 1s after the offset of test odors (2s in total). For excitation during baseline and ITI, 1mw laser was delivered just before sample odors or in the middle of ITI (5-s interval in the middle of 10s) with 5-s duration, which is the same length with excitation during delay.

#### Optogenetic inhibition and inhibition of VTA^DA^ neurons

To test the effect of suppressing VTA^DA^ neurons during delay period, 5mW green laser (532nm) was delivered in a 5-s interval in the middle of 6-s delay. In addition, 2mW blue laser was delivered for VGat-ChR2 mice for indirectly inhibition of VTA^DA^ neurons.

#### Optogenetic excitation of dopaminergic terminals

To test the effect of elevating mPFC dopamine in different phases of the delay period, laser was delivered during the early delay (first 2 sec) and late delay (last 2 sec). Consecutive for the six learning days, 8∼10 mW constant blue laser (473 nm) illumination was delivered in the early delay for 13 mice and the late delay for another 13 mice. Ten cage-mate mice with control virus injected were used control, half with early-delay delivery and the other half with late-delay delivery. Several days following learning to keep the behavior reaching well-trained phase (> 80%), laser illumination was implemented for testing the behavioral effect of dopamine elevation in well-trained phase.

To test optogenetic excitation of dopaminergic terminals during odor delivery and baseline, nineteen mice were delivered with blue laser in sample-odor, test-odor, and response delay periods in 5 consecutive learning days. For activation in baseline experiment, another group of mice (10 and 9 mice for ChR2 and control group, respectively) were trained ODPA task, with 2-sec blue laser illumination during baseline period before sample odor onset and 4 sec for the delay period.

To test elevate dopamine during the entire delay period, AAV-ChR2 mice were trained for ODPA task with either 6-or12-sec delay period. Blue laser was delivered for the middle 5 sec and 11 sec during the delay period, leaving 0.5-sec gaps after sample odor offset and before test odor onset.

#### Optogenetic inhibition of dopaminergic terminals in delay

To test suppress dopaminergic terminals during delay, 10 mW green laser (532 nm) was on in 4.5 out of 5 sec delay with 0.5 s interval following sample odor offset. During the last 0.5 sec of the delay period, the laser power was ramped down to zero.

### Fiber photometry imaging

#### Data acquisition

Calcium imaging and dopamine signal recordings were collected by a single-channel fiber photometry system (Thinker Tech Nanjing Biotech Co., Ltd.) at 100 Hz. ∼30 μW-470 nm blue LED light^61^ was used to illuminate fluorescence of calcium and DA sensor. Mice were trained on a ball blowing with air. Comparing to keep mouse sitting inside a tube, allowing mouse to free run on a ball was more effective to reduce motion artifacts and improve imaging quality. Imaging was started from the first learning day after habituation and shaping. The ‘system-baseline signals’ (F0) of the imaging system were acquired in a dark environment for about two minutes before we plugged the optical fiber to the adaptor attached into the brain. After connecting the fiber and the adaptor, we warmed up the imaging system and acquired the sensor signal for about five minutes before task training. Imaging was then continued until mice finished 200 trials of the DPA task in each session for 6 consecutive learning days. The sampling rate was set to 100 Hz and the raw data was binned by a 100-msec window.

#### Data analysis

After subtracted by system-baseline signals (F0), the acquired signals (F) were grouped according to the trial structure. To analyze the data, second-order exponential regression was used for fitting the calcium data and the MATLAB function “msbackadj” with a moving window of 20% of each trial was applied to correct the drift of fluorescence over long imaging ^62^. The instantaneous fluorescence change of dopamine signals was calculated by:

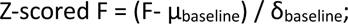

F was drift-corrected data of raw fluorescence across time. μbaseline and δbaseline was the mean and standard deviation of corrected baseline (1 s before sample odor onset), respectively.

To measure stimulus-evoked changes of dopamine release, we calculated the peri-stimulus time histogram (PSTH) of aligned trials to the onset of sample odors. The shadow was the SEM of Z-scored F cross trials (Fig.2d, Extended data Fig.3, Extended data Fig.7d).

To compare the stimulus-evoked dopamine release among different learning sessions, the peaks of task related events evoked fluorescence response and their latencies were compared in 6 learning days (Extended data Fig.2). Group differences were analyzed using Two-way ANOVA with repeated measure.

### In vivo electrophysiology

#### Electrodes array construction

Multi-electrode arrays with optic fiber were home-made as described previously^60,63^. Briefly, polyimide insulated Ni-Chrome wires (diameter:12.5μm) were twisted into tetrodes and inserted into two 2 × 4 silicon tubes (diameter 50 μm) for two-hemisphere insertion into mPFC, with 32 channels in each hemisphere. The silicon tubes were adhered to a 3D-printed resin chamber. The tetrodes outside of the tube array were adhered to a nut to move while rotating a screw. Gold planting was processed on the tips of wires before implantation to keep the impedance below 1 MΩ.

#### Experimental design

For recoding in VTA, we recorded extracellularly from multiple neurons in VTA. Before or after each behavioral session, we delivered trains of 10 laser pulses, each 5-ms long, at 1, 5, 10, 20,40 Hz at 473nm at 5-10mW for optogenetic tagging dopamine neurons. the neuron that the spike followed the laser pulse at every frequency visually the correlation coefficient was larger than 0.9 was defined optogenetic-tagged dopamine neurons.

For recording in mPFC and optogenetic perturbation on VTA^DA^ neurons, mice performed the ODPA task with blue (473nm, 1mW) or green (532nm, 5mW) laser illumination during the delay period with blocked design (20 trails were laser-on and 20 trials were laser-off, 200 trials in total in a session). To avoid excessive laser stimulation causing mice not engaging in the task, green laser was delivered unilaterally.

For recording in mPFC and optogenetic excitation of dopaminergic terminals, mice performed the ODPA task with laser delivery of a blocked design (schedule within a block: 20 early-on trials; 20 laser-off trials; 20 late-on trials). Each day mice performed 5 blocks (300 trials). 5∼8mW blue laser was delivered at 20 Hz with 5ms duration during early- or late-delay period^18^. To eliminate the performance instability of the initial trials and balance the number of laser-on trials, the first 20 and last 40 trials of all sessions were not included in data analysis. There was no significant change in behavioral performance under this design, which might be due to either experimental design (20-Hz pulsed stimulation here, vs. constant light on in Fig.3) or less coverage area of single optical fiber (vs. two fibers, one in each hemisphere in Fig.3).

Recording sites were further verified histologically with electrolytic lesions using 10s of 50 μA direct current at 1Hz and from the optical fiber track.

#### Electrophysiological Recording and spike sorting

Neural activity was continuously recorded using Plexon MPA Data Acquisition System. Wild-band signals (0.5 - 8000 Hz) were amplified (× 20000) and digitized at 40 kHz. Each day op-tetrodes were deepened 50μm after training to reduce the resampling of same neurons.

Spike sorting was preformed based on the waveforms using Plexon offline sorter software as in ref 27, 34. Briefly, spikes were clustered in the principal components analysis (PCA) space based on waveforms. Single units were sorted according to the distance in PCA space and visually verified for the temporal stability. Signal-to-noise ratio (the mean peak amplitude of all spikes divided by the standard deviation during baseline), false alarm rate (FA, the ratio of spikes within 2 ms in ISI correlogram comparing the total spike number) and firing rate were measured to qualify single units. Single units were excluded with a firing rate less than 2 Hz and FA larger than 0.15%^63^. The cross-session recorded units with a waveform correlation coefficient larger than 0.7 were regarded as same units and were excluded.

#### Data analysis

After spike sorting, all data were analyzed in MATLAB. To avoid double counting, we got rid of the units with high correlation coefficient (>90%) for waveform and firing characteristics cross sessions. To measure firing rate, peri-stimulus time histograms (PSTHs) were constructed using 100-ms bin. To quantify the neuronal modulation of all recorded units, firing rate was normalized to the baseline (1s before sample odor onset) as normalized firing rate. PSTHs and heatmaps plot the trial-averaged firing rate with a 500-msec bin and 100-msec step size.

To analyze the odor selectivity, various trials were grouped according to the identity of odors. We calculated the selectivity using 2-sec bin during the delay period. Neuronal selectivity was computed as the formula: Selectivity = (FRS1 – FRS2) / (FRS1 + FRS2) (Fig.1k). We defined a statistically significant neuron in any second of the delay period as a selective neuron (Fig.1l, Extended data Fig.4d).

To quantify the modification on neuronal activity by optogenetic illumination, we defined modification index to measure the laser effect which calculated as the raw firing rate within 500ms during laser illumination normalized to the firing rate in the baseline period.

#### MCC decoding and Cross-temporal decoding analysis

To assess the population coding ability of neurons in VTA for sample odors, we performed population decoding analysis (Extended data. 4f) by using the maximum correlation coefficient (MCC) classifier with the neural decoding tool box^63,64^. All selective neurons were included in this analysis. For MCC decoding, the firing for each trial was binned in 500ms with 100ms step for each neuron. Firing of one and the left trials was used as test and training data. to obtain the decoding accuracy, this procedure was repeated for 500 times. To quantify the persistence of decoding during delay, we performed the cross-temporal decoding (CTD) analysis by using MCC classifier (Fig.1m). We trained the classifier with specific time bin and test the classifier for all the time bin during the whole trial. Repeating this process for every time bin produced a cross temporal decoding matrix. The decoding analysis was repeated for 500 times.

#### SVM decoding

Liner support vector machines (SVM) were used to analyze the population decoding in MATLAB by the function *“fitcsvm”*. The normalized spike counts during early- or late-delay period for each neuron in each trial were used. For each repeat, all trials labelled with sample odor identity were randomly partitioned to a training set and a test set in a ratio 9 to 1. Form the training set, the normalized firing of the selected neural population was used to train the SVM with the liner function kernel. From the test set, each trial was used to test the classification accuracy. All trials were then randomly re-partitioned and averaged decoding accuracy was obtained with 1000 repeats. For the shuffled control, the procedure was repeated 1000 times with the nominal sample odor for each trial randomly redistributed (Fig.1s,y, Fig.4j,k, Extended data Fig.5r,u, Extended data Fig.6r,u).

To analyze cell type, all units were further classified as broad-spiking (BS) putative pyramidal neurons and narrow-spiking (NS) putative interneurons based on the distribution of the trough-to-peak distance^65–67^. A unit was classified as BS neuron if its trough-to-peak distance was > 350μs or as a NS neuron if its trough-to-peak distance was ≤ 350μs in this paper^68^. (Fig.4i, Extended data Fig.10b,e).

### Histology

#### Perfusion and tissue storage

Mice were deeply anaesthetized with sodium pentobarbital (120 mg/kg; intraperitoneal injection) and transcardially perfused with 20 mL normal PBS followed by 20 mL of cold 4 % paraformaldehyde (PFA) dissolved in PBS. After decapitation, the brain was extracted from the cranial cavity and placed in 4 % PFA solution and stored at 4 ℃ for 24 h (for DPAI staining) or 4 ∼ 6 h (for other staining), then transferred to PBS. Brains were sectioned at 80 μm by vibratome (Leica) coronally. Sections were stored in PBS at 4 ℃ until immunohistochemical processing.

#### Immunohistochemistry

For DAPI staining, brain sections were incubated in DAPI solution (C1002, Beyotime, 1:1000 diluted in PBS) for 10 min, then consecutively rinsed in PBS for 3 times (10 min each time). For tyrosine hydroxylase (TH) staining, brain sections were firstly treated in the blocking solution (0.3% Triton + 3% BSA in PBS, Jackson ImmunoResearch) for 1 h at home temperature, then followed by incubating overnight at 4 ℃ in the primary antibody solution: mouse anti-TH (1:1000; Jackson ImmunoResearch) in antibody diluent (0.15% Triton and 1.5% BSA in 1 × PBS). Sections were then washed 3 times in PBS and immediately transferred to the secondary antibody solution: Cy3 goat anti-mouse (1:500, Jackson ImmunoResearch) for 2 hours at home temperature. Then, sessions were washed 4 times in PBS (10 min each time) and DPAI (1:1000) diluted in PBS solution and mounted onto glass slides. Sections were allowed to dry and were coverslipped using 50% glycerinum in PBS.

#### Microscopy

Fluorescence images were obtained with Olympus VS120 using a 10X (NA = 0.45) lens. Images were analyzed with OlyVIA (Olympus).

### Blind design

For all optogenetic experiments, mouse ID for the control and experimental groups were labelled by other people in the lab. The group information was not available to the experimenter until the end of experiments.

### Statistics

Group comparisons cross days were two-way ANOVA with repeated measure. Wilcoxon rank Sum test was used for performance within session. Permutation test was used for selectivity of single cell, with 1000 times resampling from 40 trials.

## QUANTIFICATION AND STATISTICAL ANALYSIS

## DATA AND CODE AVAILABILITY

## Acknowledgements

We thank Drs. Muming Poo and Yang Dan for their comments on the manuscript. The work was supported by National Key R&D Program of China (Grant No. 2019YFA0709504), the Innovations of Science and Technology 2030 from the Ministry of Science and Technology of China (Grant No. 2021ZD0203601), National Natural Science Foundation of China (Grant No. 31827803 and 32161133024), the Strategic Priority Research Program of the Chinese Academy of Sciences (Grant No. XDA27010400 and XDB32010100), the Shanghai Municipal Science and Technology Major Project (Grant No. 2018SHZDZX05 and 2021SHZDZX), the Lingang Laboratory (Grant No. LG202105-01-01), Shanghai Pilot Program for Basic Research-Chinese Academy of Science, Shanghai Branch (JCYJ-SHFY-2022-010).

## Author contributions

C.F.G. and C.Y.L. designed the study. C.F.G. conducted the optogenetics and fiber photometry imaging, extracellular recording, injection, and histology and analyzed these data. Z.Q.C provided help in virus injection and histology. H.M.F helped for construction of all electrodes. R.Q.H provided help in virus injection. F.M.S. and Y.L.L. provided the virus of dopamine sensor and helped in interpreting analysis results for sensor imaging. C.F.G. and C.Y.L. interpreted the data and wrote the manuscript.

