## Supplementary material for "Timing-dependent modulation of working memory by VTA dopamine release in medial prefrontal cortex": Ge et al. Figure

### Extended Data Fig.1| Behavioral performance during learning.

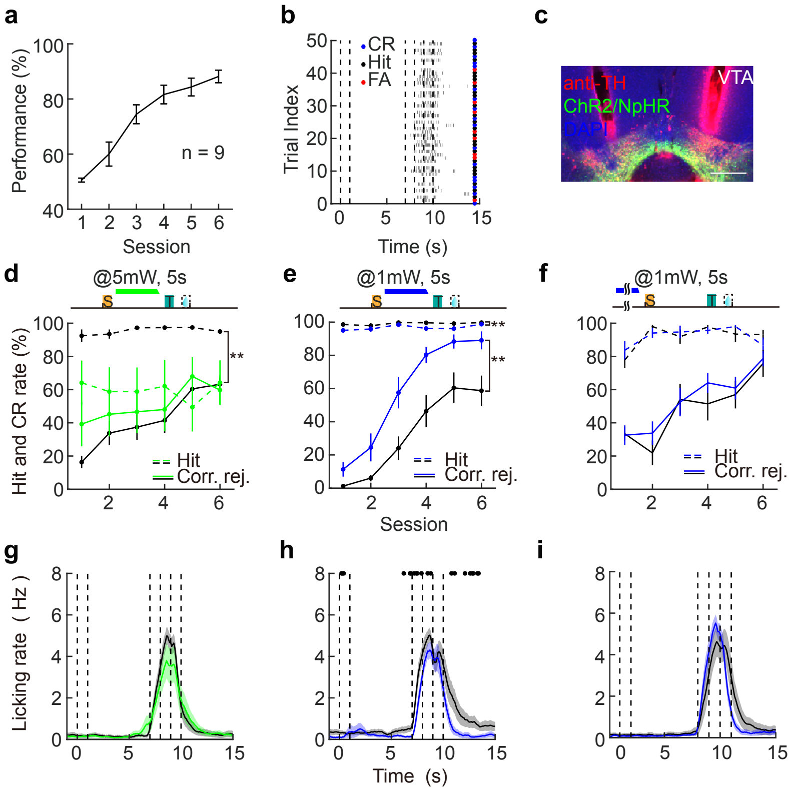

**Extended Data Fig.2| Behavioral results for optogenetic excitation of VTADA and VTAGABA neurons in learning the ODPA task.**

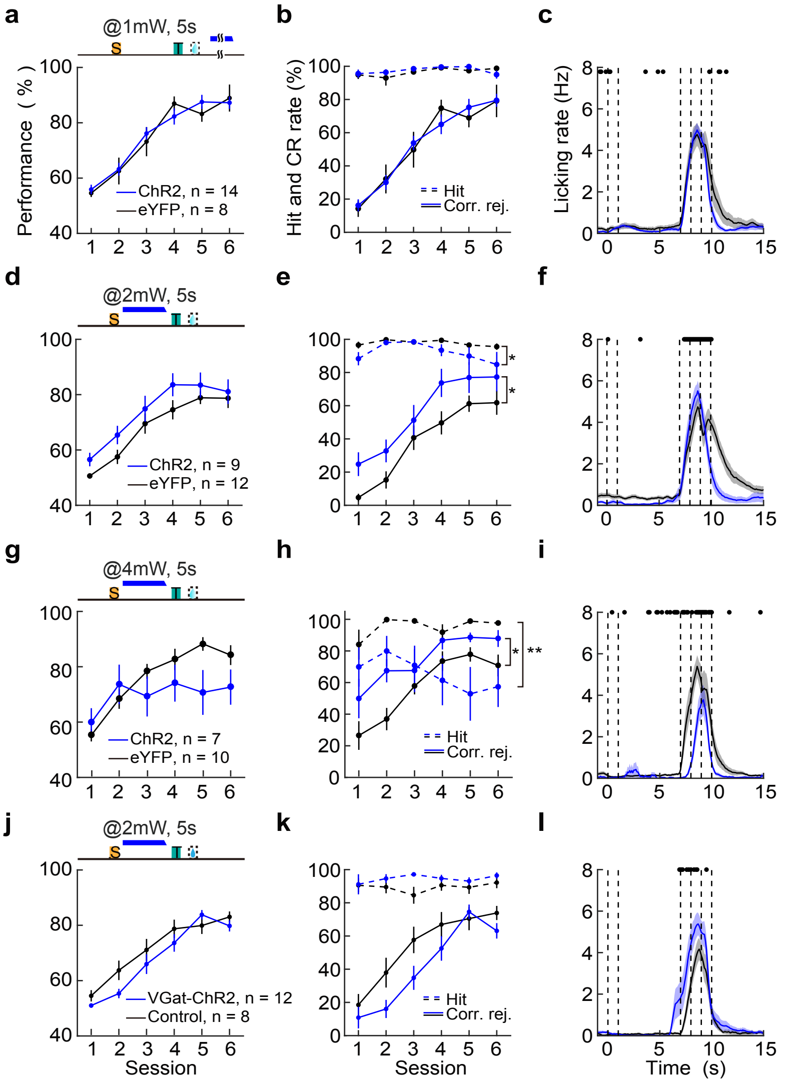

**Extended Data Fig. 3| Population calcium signals of VTADA neurons in OPDA task.**

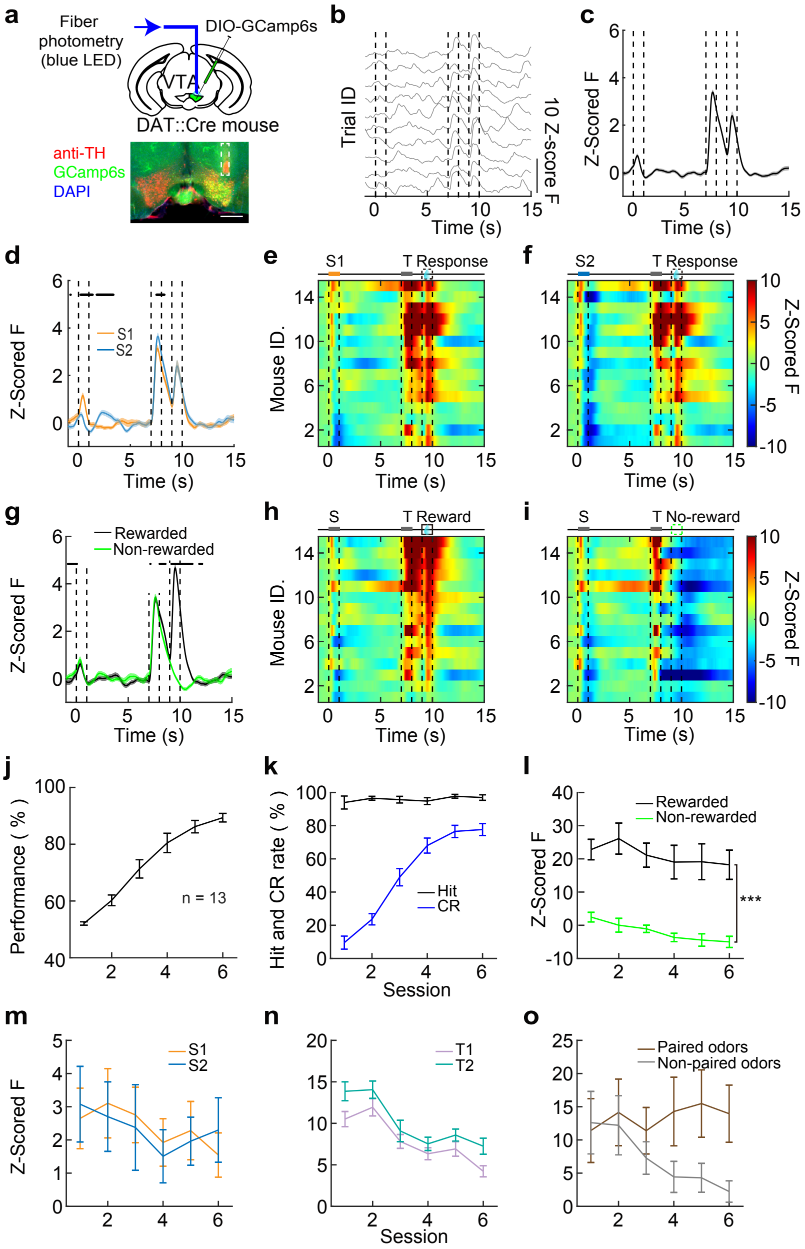

**Extended Data Fig.4| Electrophysiological recoding in VTA.**

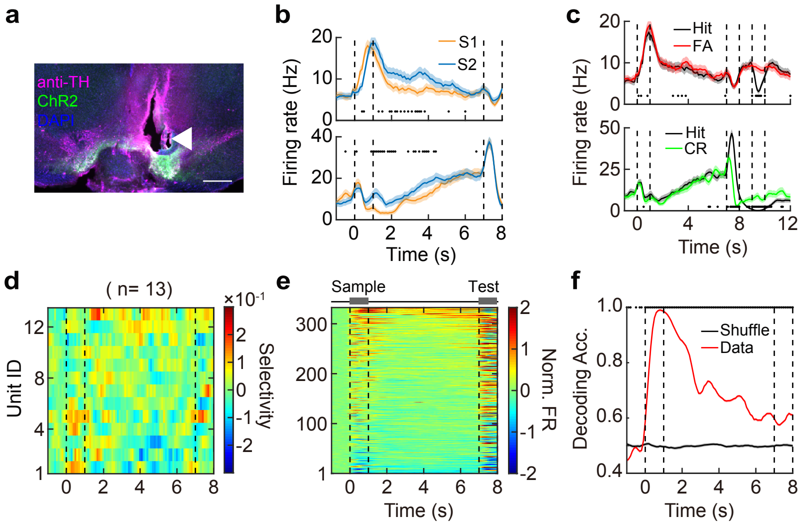

**Extended Data Fig. 5| Electrophysiological recoding in mPFC with optogenetic excitation of VTADA neurons during the delay per**

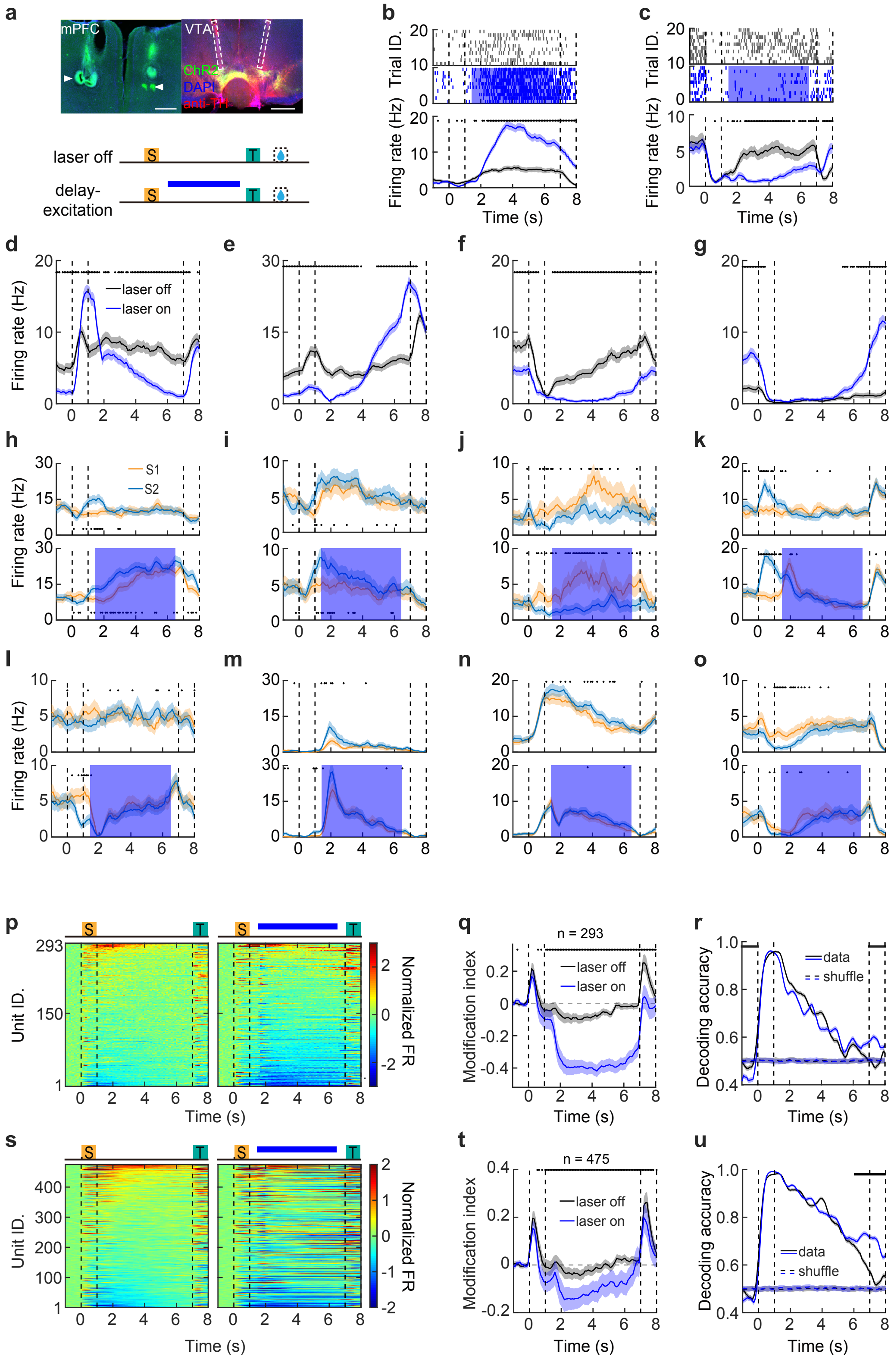

**Extended Data Fig. 6| Electrophysiological recoding in mPFC with optogenetic suppression of VTADA neurons during the delay p**

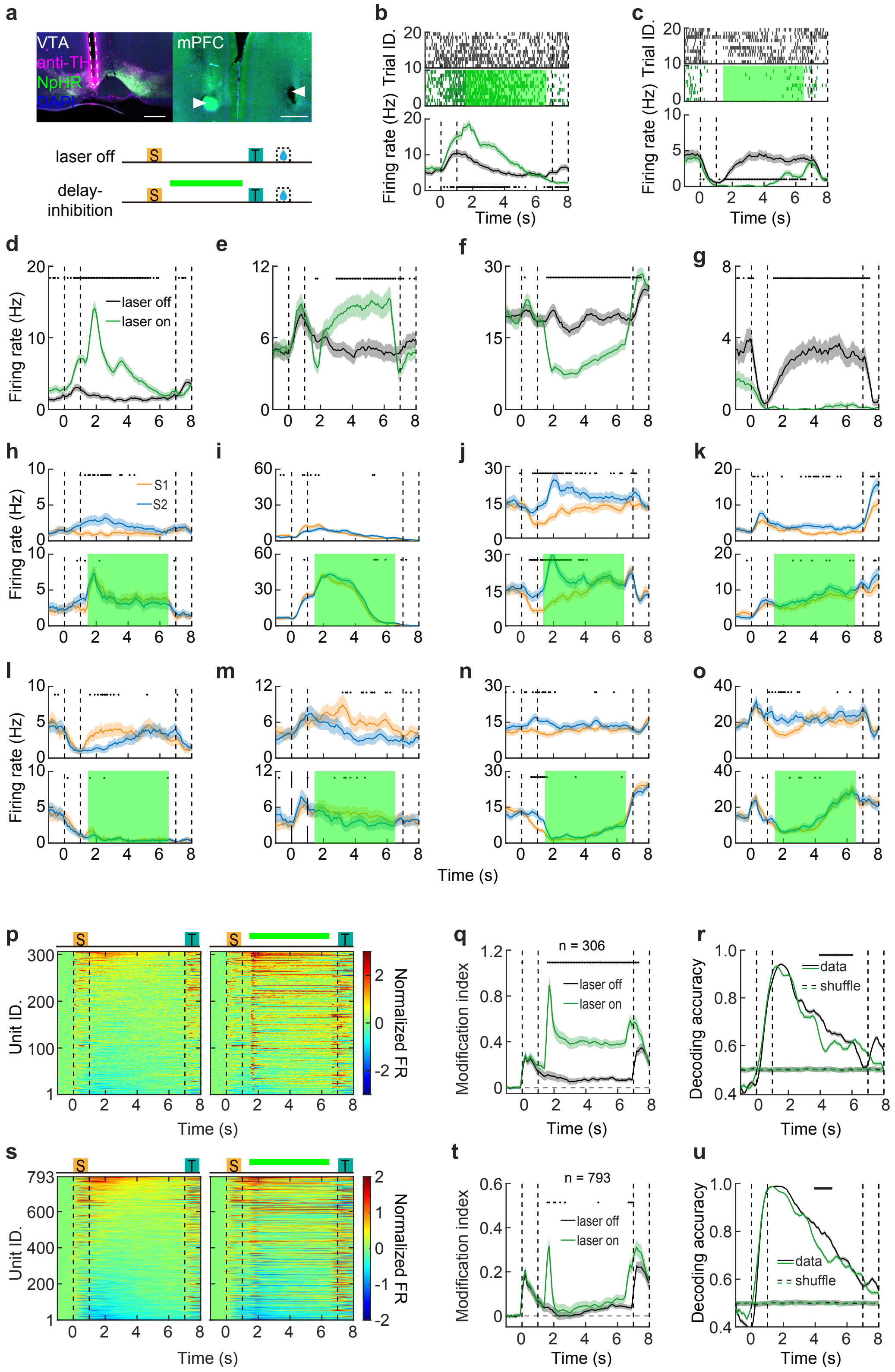

**Extended Data Fig.7| Behavioral performance and dopamine response in mPFC during learning.**

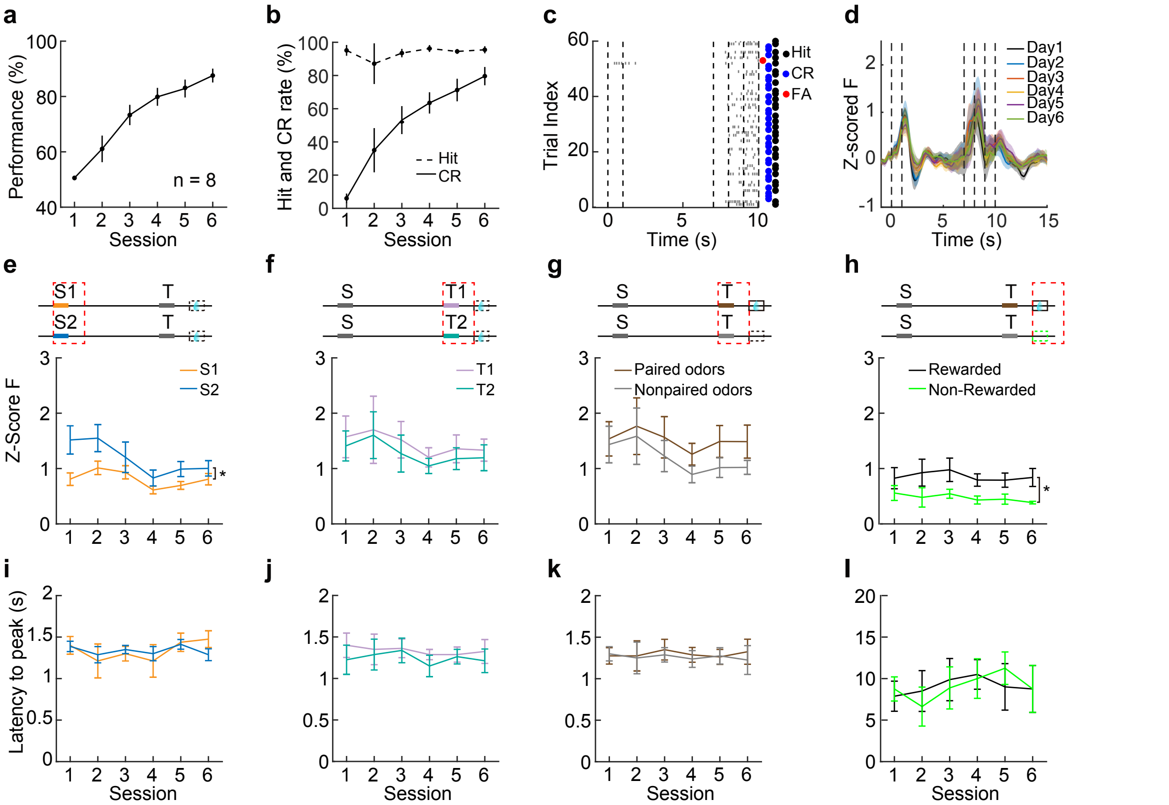

**Extended Data Fig.8| Behavioral performance of optogenetic manipulation on VTA dopaminergic terminals in mPFC.**

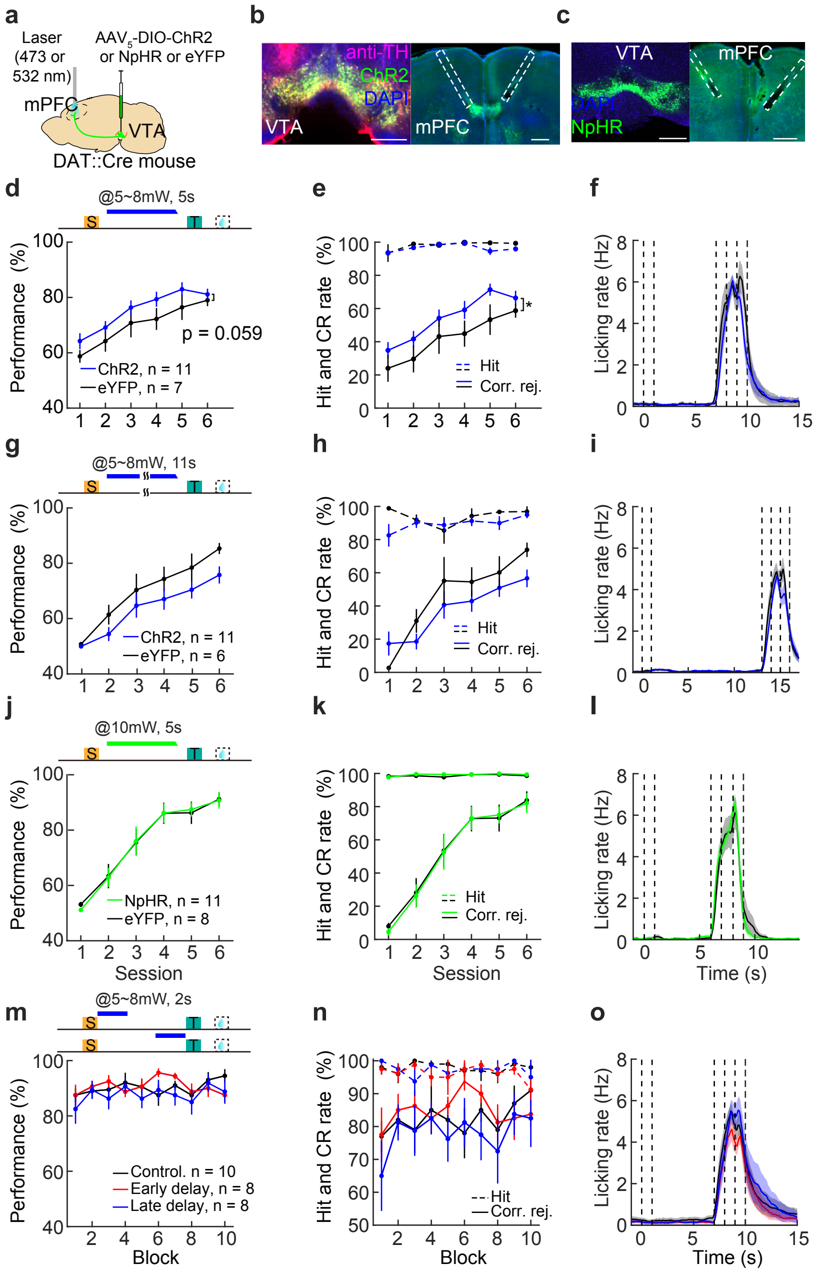

**Extended Data Fig. 9| Neural modulation of mPFC neurons induced by optogenetic excitation of dopaminergic terminals.**

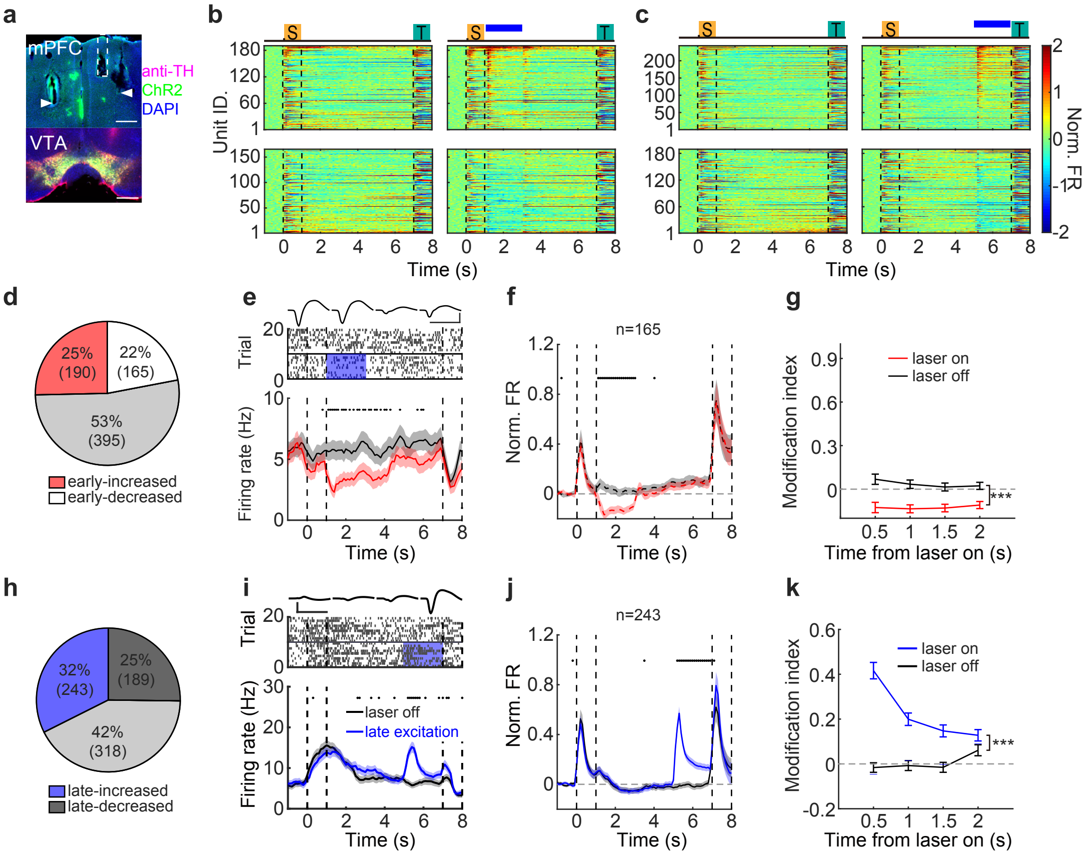

**Extended data Fig.10| Modulation in memory-coding ability of mPFC neurons by optogenetic excitation of dopaminergic terminal**

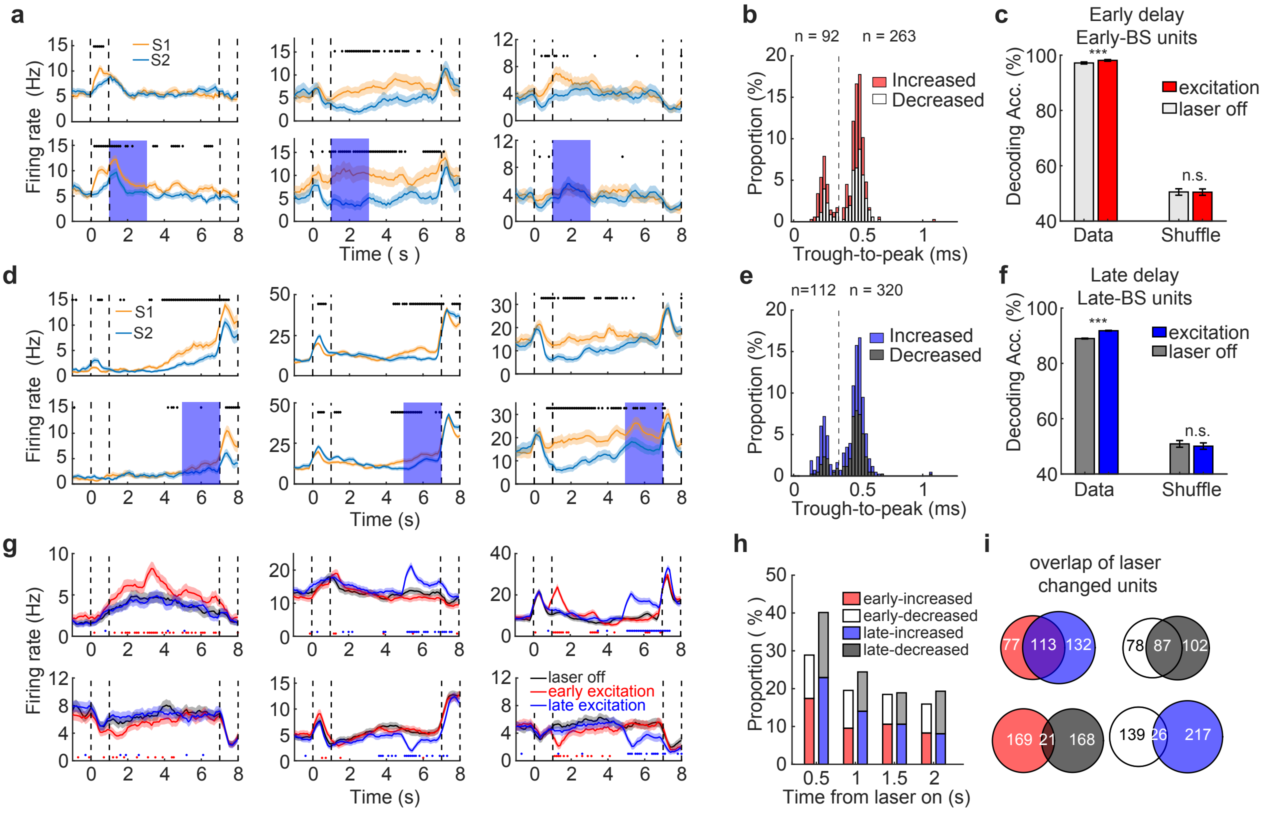
